# Genome-wide landscape of m^6^A regulatory genes in *Marchantia polymorpha*: conservation, diversification, and expression dynamics

**DOI:** 10.64898/2026.09.29.755302

**Authors:** Joong-Tak Yoon, Shaban Muhammad, Yicheng Liu, Eun Yu Kim

**Affiliations:** Division of Natural and Applied Sciences, Duke Kunshan University, Kunshan, Jiangsu 215316, China; Environmental Research Center, Duke Kunshan University, Kunshan, Jiangsu 215300, China; Center for Genome Engineering, Institute for Basic Science, Daejeon 34126, Republic of Korea

**Author notes:** These authors contributed equally to this work (Joong-Tak Yoon, Shaban Muhammad). Correspondence: Joong-Tak Yoon, Eun Yu Kim.

## Abstract

**Background:** N^6^-Methyladenosine (m^6^A) regulates RNA fate in flowering plants, but the repertoire and expression patterns of m^6^A-related genes remain incompletely characterized in liverworts. Here, we characterized these gene families in the liverwort *Marchantia polymorpha* within a comparative framework spanning diverse plant lineages.

**Results:** A comparative survey of 119 plant genomes using sequence similarity and conserved-domain criteria identified 18 candidate m^6^A-related loci in *M. polymorpha*, comprising seven writer-related genes, five AlkB homolog genes, and six YTH- or CPSF30-related genes. Across the surveyed genomes, canonical writer components generally had low copy numbers, whereas AlkB homolog and reader-related families showed greater copy-number variation. Phylogenetic and domain analyses supported the family assignments of the *M. polymorpha* candidates. Eight of the 18 candidate loci showed detectable collinearity with *Marchantia quadrata*. Putative promoter analysis identified motifs annotated as hormone-, light-, and stress-responsive, with hormone-responsive motifs accounting for the largest share of predicted sites. Public transcriptome analysis revealed variation in candidate gene expression across developmental stages and tissues. Quantitative reverse-transcription PCR showed higher transcript abundance for all six YTH- or CPSF30-related genes in thalli than in gemmae or gemma cups. Cold, heat, osmotic, salt, and dark treatments elicited gene-specific expression responses. Darkness increased transcript abundance of several candidates, including *MpMTA*, *MpALKBH1A*, and *MpECT2*. A predicted protein-association network suggested links between subsets of the candidates and RNA-processing proteins.

**Conclusions:** These findings define a candidate m^6^A-related gene repertoire in *M. polymorpha* and identify tissue- and stress-responsive genes for targeted functional studies of RNA methylation in this liverwort.

## Background

N^6^-Methyladenosine (m^6^A) is a widespread internal RNA modification that adds a post-transcriptional layer to gene regulation (Luo et al., 2014; Zhong et al., 2008). On plant transcripts, m^6^A has been associated with RNA processing, alternative polyadenylation, transcript stability, and translation (Chen et al., 2024; Hou et al., 2021; Parker et al., 2022; Song et al., 2023). Its effects depend on the modified RNA and the proteins that install, remove, or recognize the methyl group (Duan et al., 2017; Ruzicka et al., 2017; Song et al., 2021). These proteins are commonly classified as writers, erasers, and readers, respectively (Figure 1A) (Duan et al., 2017; Ruzicka et al., 2017; Song et al., 2021). Studies in flowering plants have linked this regulatory framework to embryogenesis, organ development, and responses to environmental signals, but the functions of individual family members can differ across biological contexts (Cai et al., 2024; Ruzicka et al., 2017; Shen et al., 2016; Zhong et al., 2008).

**Figure 1.**
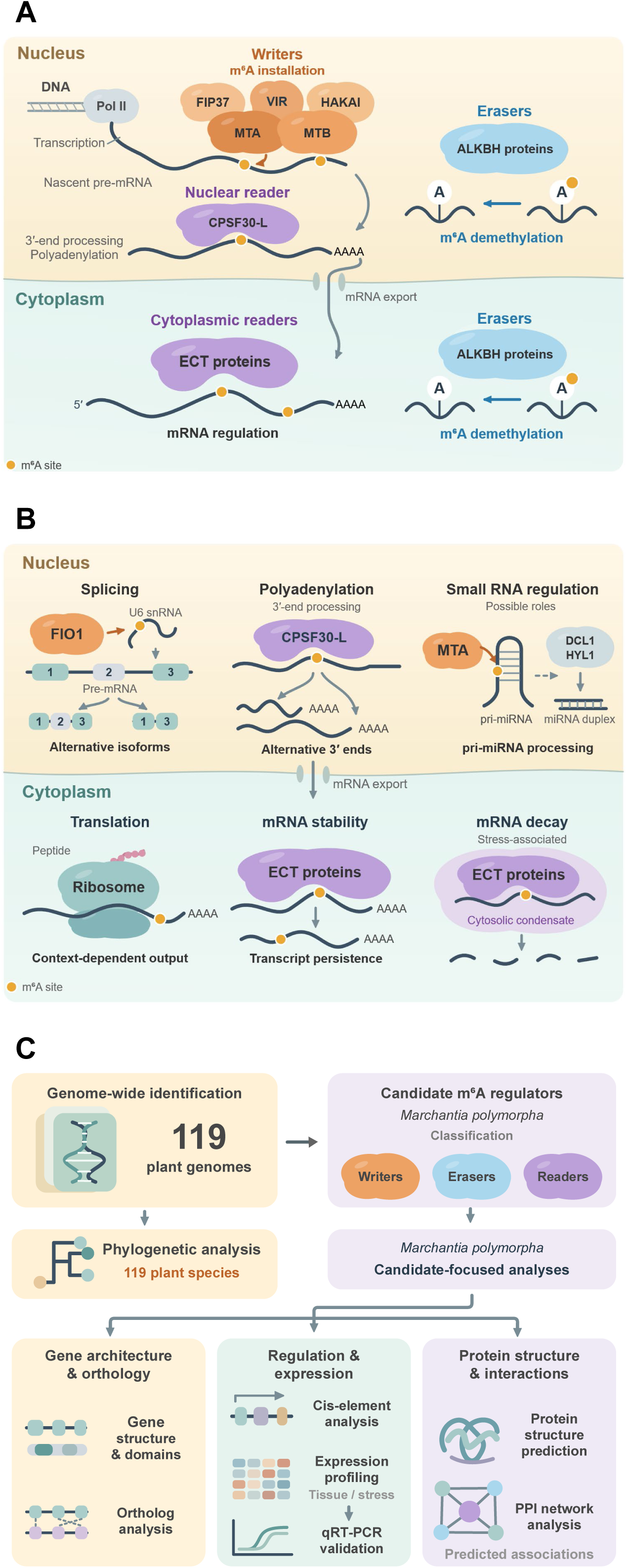
Plant m^6^A regulatory pathways and the analytical workflow used in this study. **a** Schematic of m^6^A regulation in plants. In the nucleus, RNA polymerase II (Pol II) transcribes DNA into precursor mRNA (pre-mRNA), and the writer complex containing MTA, MTB, FIP37, VIR, and HAKAI adds m^6^A to RNA. The long isoform of cleavage and polyadenylation specificity factor 30 (CPSF30-L) is shown as a nuclear reader associated with 3′-end processing and polyadenylation. Following mRNA export, evolutionarily conserved C-terminal region (ECT) proteins are shown as cytoplasmic readers. Selected AlkB homolog (ALKBH) proteins are shown as nuclear or cytoplasmic erasers. Orange protein shapes denote writers, lavender shapes readers, and pale-blue shapes ALKBH proteins; yellow circles on RNA denote N^6^-methyladenosine (m^6^A), whereas white circles labeled A denote unmodified adenosine. The one asterisk after ALKBH proteins indicates that the diagram illustrates a general mechanism of m^6^A demethylation by ALKBH-family erasers. The horizontal divider separates the nucleus from the cytoplasm; arrows indicate the illustrated RNA-processing and transport steps. **b** Examples of plant RNA processes associated with the proteins in **a**. FIONA1 (FIO1) is depicted alongside U6 small nuclear RNA (snRNA) and pre-mRNA splicing; CPSF30-L alongside alternative 3′-end formation; MTA alongside primary microRNA (pri-miRNA) processing; and ECT proteins alongside translation, transcript stability, and stress-associated mRNA decay in cytosolic condensates. Yellow circles represent m^6^A sites; the pink chain emerging from the ribosome represents the translation product. DICER-LIKE 1 (DCL1) and HYPONASTIC LEAVES 1 (HYL1) are indicated in the small-RNA pathway. These schematics do not establish the corresponding functions in *Marchantia polymorpha*. **c** Workflow of the present study. Genome-wide searches across 119 plant genomes identified candidate m^6^A regulators; the *M. polymorpha* candidates were classified as writers (orange), erasers (light blue), or readers (lavender). Subsequent analyses examined phylogenetic relationships, gene structure, conserved domains, chromosomal collinearity, *cis*-elements, expression profiles, tissue- and stress-response quantitative reverse-transcription PCR (qRT-PCR) results, predicted protein structures, and a protein-protein interaction (PPI) network of predicted associations. Pale orange, mint, and lavender background panels group the gene architecture/ collinearity, regulation/expression, and protein structure/interaction analyses, respectively; grey arrows indicate the analysis sequence.

In *Arabidopsis thaliana*, METHYLTRANSFERASE A (MTA) and METHYLTRANSFERASE B (MTB) are central components of the mRNA m^6^A writer complex, together with FKBP12 INTERACTING PROTEIN 37 KD (FIP37), VIRILIZER (VIR), and HAKAI (Ruzicka et al., 2017; Zhong et al., 2008). Depletion of several of these components lowers mRNA methylation and can disrupt embryogenesis, shoot meristem activity, and root development (Ruzicka et al., 2017; Shen et al., 2016; Zhong et al., 2008). The effects of the writer machinery also extend beyond protein-coding mRNAs (Bhat et al., 2020). MTA-dependent methylation of primary microRNAs (pri-miRNAs) promotes their efficient processing by the nuclear microRNA machinery, which includes DICER-LIKE 1 (DCL1) and HYPONASTIC LEAVES 1 (HYL1) (Figure 1B) (Bhat et al., 2020). FIONA1 (FIO1), a methyltransferase related to METHYLTRANSFERASE-LIKE 16 (METTL16), represents a distinct route to RNA methylation: its modification of U6 small nuclear RNA (snRNA) affects splice-site selection and pre-mRNA splicing (Parker et al., 2022). Thus, the presence of a methyltransferase-related domain does not necessarily imply membership in the MTA-MTB writer complex or the same RNA substrate specificity (Parker et al., 2022; Ruzicka et al., 2017).

Removal of m^6^A adds another potential level of regulation (Duan et al., 2017; Martinez-Perez et al., 2017). The *A. thaliana* AlkB homolog (ALKBH) protein ALKBH10B demethylates transcripts encoding flowering regulators, including FLOWERING LOCUS T (FT), SQUAMOSA PROMOTER BINDING PROTEIN-LIKE 3 (SPL3), and SQUAMOSA PROMOTER BINDING PROTEIN-LIKE 9 (SPL9), and thereby affects their stability and the timing of floral transition (Duan et al., 2017). A different family member, ALKBH9B, influences the m^6^A content of alfalfa mosaic virus RNA and viral infection (Martinez-Perez et al., 2017). These examples show that plant ALKBH proteins can act in different cellular and biological contexts (Figure 1A) (Duan et al., 2017; Martinez-Perez et al., 2017). However, RNA demethylase activity and substrate choice have been demonstrated for only a subset of plant ALKBH proteins; they cannot be inferred for every family member from sequence similarity alone (Duan et al., 2017; Martinez-Perez et al., 2017; Zhang et al., 2024).

The consequences of m^6^A are also determined by reader proteins (Song et al., 2023; Song et al., 2021). In *A. thaliana*, the long isoform of CLEAVAGE AND POLYADENYLATION SPECIFICITY FACTOR 30 (CPSF30-L) contains an m^6^A-recognizing YT521-B homology (YTH) domain and contributes to alternative polyadenylation of target transcripts in the nucleus, with reported effects on nitrate signaling, floral transition, and abscisic acid (ABA) responses (Hou et al., 2021; Song et al., 2021). In the cytoplasm, EVOLUTIONARILY CONSERVED C-TERMINAL REGION (ECT) proteins constitute another group of YTH-domain readers (Arribas-Hernandez et al., 2018; Song et al., 2023). *A. thaliana* ECT2, ECT3, and ECT4 support organogenesis and can promote the stability of bound transcripts; they have also been implicated in translation of immune-related mRNAs under specific conditions (Arribas-Hernandez et al., 2018; Arribas-Hernandez et al., 2020; Chen et al., 2024; Song et al., 2023). ECT8 provides a distinct stress-associated example: it limits translation by sequestering m^6^A-modified transcripts in stress granules during ABA responses, whereas its association with processing bodies promotes target mRNA decay in the context of salt stress (Cai et al., 2024; Wu et al., 2024). Accordingly, reader-dependent outcomes range from RNA persistence to selective translation or turnover and should be evaluated in the relevant cellular context (Figure 1B) (Cai et al., 2024; Chen et al., 2024; Song et al., 2023; Song et al., 2021; Wu et al., 2024).

Genome-wide and transcriptome-scale studies in *A. thaliana*, *Oryza sativa*, *Litchi chinensis*, *Pinus massoniana*, poplar, and *Brassica napus* have catalogued m^6^A-related gene families and described variation in copy number, promoter motifs, and developmental or stress-associated expression (Hasan et al., 2024; Nishat et al., 2022; Shan et al., 2025; Sun et al., 2022; Tang et al., 2022; Yao et al., 2024). A broader comparative analysis across 154 plant species examined the evolutionary relationships, family expansions, and conserved genomic contexts of m^6^A-related proteins (Zhang et al., 2024). The liverwort *Marchantia polymorpha* offers a useful comparison with flowering plants because its life cycle includes distinct vegetative, asexual, and sexual developmental contexts (Bowman et al., 2017). Homologs of the core writer-complex components MTA, MTB, FIP37, VIR, and HAKAI have already been identified in *M. polymorpha (Pietrykowska et al., 2022)*. In addition, a *M. polymorpha* YTH-domain family (YTHDF) protein partially complemented leaf-development defects in *A. thaliana ect2 ect3 ect4* mutants in a heterologous assay (Flores-Tellez et al., 2023). These findings establish a basis for investigating m^6^A-related proteins in *M. polymorpha*, but do not resolve how the broader candidate repertoire is expressed across developmental contexts and abiotic stress conditions. Integrating comparative sequence analyses with expression profiling can help identify candidates for functional investigation.

Here, we surveyed candidate m^6^A-related proteins across 119 plant genomes and identified 18 candidate loci in *M. polymorpha*: seven writer-related genes, five ALKBH-family genes, and six YTH- or CPSF30-related genes (Figure 1C). We combined phylogenetic and domain analyses with gene and predicted protein structures, chromosomal collinearity with *M. quadrata*, putative promoter motifs, expression across developmental contexts and abiotic stresses, and predicted protein associations. This integrated analysis defines a candidate repertoire and identifies specific genes for future tests of RNA methylation and function in a liverwort.

## Methods

### Plant materials and growth conditions

The male *Marchantia polymorpha* accession Takaragaike-1 (Tak-1) was propagated on half-strength Gamborg’s B5 medium with vitamins (Duchefa Biochemie, G0210) supplemented with 1% (w/v) sucrose (Adamas life, 66841U) and 1.4% (w/v) phyto agar (Duchefa Biochemie, P1003), adjusted to pH 5.5. Plants were grown at 22 °C under continuous LED illumination at a photosynthetic photon flux density of 100 μmol m^-2^ s^-1^.

### Abiotic stress treatments

Abiotic stress treatments were performed as previously described, with modifications (Tan et al., 2023). For salt and osmotic stress treatments, gemmae were grown for 15 days on sucrose-free half-strength Gamborg’s B5 medium with vitamins containing 1.4% (w/v) phyto agar and adjusted to pH 5.5. The medium was supplemented with 40 mM NaCl (Macklin, S805286) for salt stress or 100 mM mannitol (Greagent, G77529B) for osmotic stress. For cold, heat, and dark stress treatments, gemmae were initially grown on the same sucrose-free basal medium without NaCl or mannitol. On day 14, plates were transferred to 3 °C or 33 °C for 24 h for cold and heat treatments, respectively. For dark treatment, plates were wrapped in aluminum foil on day 12 and maintained in darkness for 3 days. Corresponding control plants were grown on the same sucrose-free basal medium without stress treatment at 22 °C under continuous LED light until day 15. On day 15, control and stress-treated plants were harvested and used for further analysis.

### Genome-wide identification of m^6^A writers, erasers, and readers in 119 plant genomes

Genome assemblies, gene annotations, and predicted protein sequences for 118 plant species were obtained from Ensembl Plants release 62 (Howe et al., 2020). The *M. polymorpha* MpTak_v7.1 assembly, annotation, and predicted protein sequences were obtained from marchantia.info, bringing the dataset to 119 plant genomes (Tanizawa et al., 2026). Previously reported *Arabidopsis thaliana* m^6^A-related proteins were used as reference queries (Hu et al., 2019; Nishat et al., 2022). The query set comprised writer-related proteins, ALKBH-family proteins, and reader-related proteins. AtALKBH1B and AtALKBH1C were excluded from the ALKBH reference query set because a functional analysis of the four *A. thaliana* ALKBH1 homologs identified AtALKBH1B as a likely pseudogene and found very low in vitro DNA N^6^-methyladenine demethylase activity for AtALKBH1C (Li et al., 2024). AtALKBH8 was excluded because its characterized function is tRNA wobble-uridine modification (Leihne et al., 2011). The names and locus identifiers of the included reference queries are provided in Supplementary Table S1.

Predicted protein sequences from the 119 species were combined into a single FASTA file, with sequence identifiers linked to their species of origin and original protein identifiers. A local protein database was constructed from the combined sequences using makeblastdb (NCBI BLAST+ v2.17.0+). BLASTp v2.17.0+ was used to search this database with the *A. thaliana* reference sequences at an E-value cutoff of 1 X 10^-5^ (Camacho et al., 2009). Hits with less than 30% amino acid identity across the aligned region were excluded. Proteins passing these criteria were extracted for conserved-domain analysis.

Candidate proteins were searched against the NCBI Conserved Domain Database (CDD v3.21) using RPS-BLAST v2.17.0+ at an E-value cutoff of 1 X 10^-3^ (Camacho et al., 2009; Lu et al., 2020). The diagnostic domain models examined were MT-A70 (CDD: 368270, pfam05063), Wtap (CDD: 465345, pfam17098), VIR_N (CDD: 464938, pfam15912), Methyltransf_10, MTase-10 (CDD: 399160, pfam05971), RING-HC_HAKAI-like (CDD: 438171, cd16508), 2OG-FeII_Oxy_2, ALKBH (CDD: 433285, pfam13532), YTH (CDD: 410979, cd21134, CDD: 461195, pfam04146), Cleavage and polyadenylation specificity factor, CPSF (CDD: 227416, COG5084), and RRM_ist3_like, RRM (CDD: 409845, cd12411). Candidates without a diagnostic domain consistent with their proposed family were excluded. The remaining proteins were classified as writer-, eraser-, or reader-related candidates based on their similarity to the reference queries and their conserved-domain composition.

For cross-species copy-number analysis, the longest protein isoform among those passing both the BLASTp and RPS-BLAST criteria was retained for each locus, and each locus was counted once within its assigned family. The same representative-isoform approach was used for phylogenetic analysis of the other species; for *M. polymorpha*, proteins annotated with the .1 isoform suffix in MarpolBase were used for phylogenetic analysis. All annotated protein isoforms of the identified *M. polymorpha* loci were evaluated for isoform-level characterization. The short MpECT12 isoform Mp3g00420.2 was excluded from downstream analyses because it lacked the YTH-domain region, leaving 23 selected protein isoforms. For the subsequent isoform-level analyses, the 23 selected transcript/protein isoforms listed in Table 1 were used.

**Table 1.**
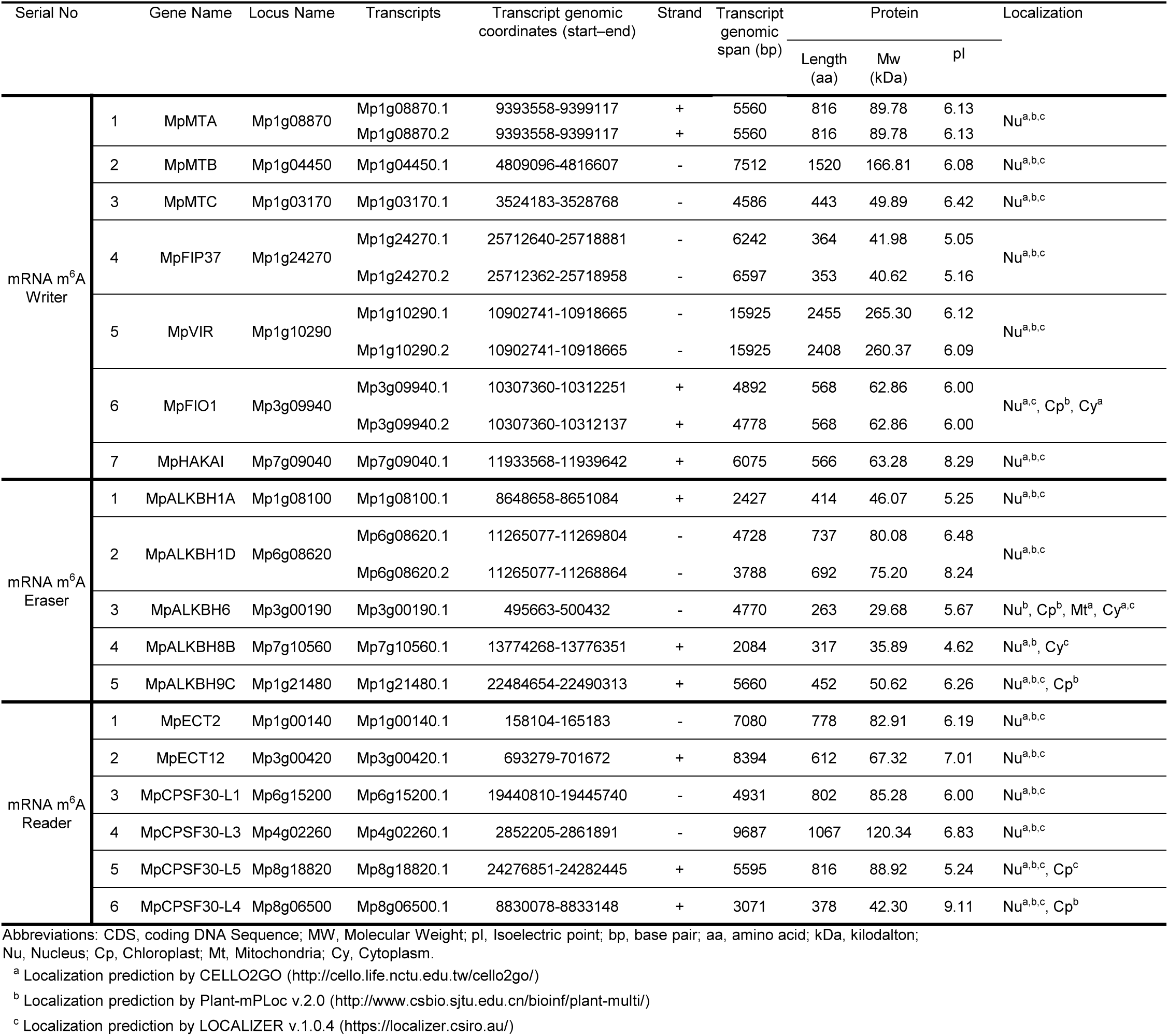
Identified candidate m^6^A-related genes in *M. polymorpha*. The table lists 18 loci - seven writers, five erasers, and six readers - and their 23 selected transcript entries. Gene names, locus and transcript identifiers, transcript genomic coordinates, strand, transcript genomic span, predicted protein length, molecular weight (MW), isoelectric point (pI), and predicted subcellular localization are shown. Localization abbreviations are Nu, nucleus; Cp, chloroplast; Mt, mitochondrion; and Cy, cytoplasm. Superscripts a, b, and c identify predictions from CELLO2GO, Plant-mPLoc v2.0, and LOCALIZER v1.0.4, respectively; predictions from different programs are reported separately. bp, base pairs; aa, amino acids; kDa, kilodaltons; Mp, *M. polymorpha*.

The theoretical molecular weights and isoelectric points of the identified *M. polymorpha* proteins were calculated using ExPASy Compute pI/Mw (Bjellqvist et al., 1994; Bjellqvist et al., 1993; Wilkins et al., 1999). Subcellular localization was predicted using CELLO2GO, Plant-mPLoc v2.0, and LOCALIZER v1.0.4 (Chou & Shen, 2010; Sperschneider et al., 2017; Yu et al., 2014). The identified *M. polymorpha* loci, selected transcript identifiers, physicochemical properties, and predicted localizations are listed in Table 1.

### Multiple sequence alignment and phylogenetic analysis

For the cross-species analyses, records sharing the same species-specific gene identifier within a conserved-domain dataset were reduced to one representative sequence. The longest sequence was prioritized, with conserved-domain and BLAST scores used to resolve ties. Full-length proteins were analyzed in five writer groups (MT-A70, Wtap, VIR_N, MTase-10, RING-HC_HAKAI-like), one eraser group (ALKBH), and four reader groups (YTH, YTH+CPSF, CPSF, and RRM). Each group was aligned independently with MAFFT v7.455 using the L-INS-i strategy, implemented with the --localpair --maxiterate 1000 options (Katoh & Standley, 2013). No trimming was applied to the resulting alignments.

A maximum-likelihood tree was inferred independently for each alignment with IQ-TREE v3.0.1 (Wong et al., 2026). Model selection was performed using ModelFinder with the -m MFP option, and branch support was assessed with 3,000 ultrafast bootstrap replicates and bootstrap-tree refinement with the -bb 3000 -bnni options (Hoang et al., 2018; Kalyaanamoorthy et al., 2017). To display the independently inferred writer trees in one panel, their consensus trees were grafted onto a predefined synthetic backbone using a custom Python script based on ETE3 (Huerta-Cepas et al., 2016). The four reader trees were assembled in the same way. The ALKBH eraser tree was displayed directly. Relationships connecting different writer or reader groups were specified by the synthetic backbones and were not inferred or assigned bootstrap support by IQ-TREE. The trees were visualized and annotated with iTOL v7 (Letunic & Bork, 2024). Group annotations were based on conserved-domain composition and similarity to *A. thaliana* reference proteins.

For analyses restricted to the identified *M. polymorpha* m^6^A-related proteins, separate maximum-likelihood trees were generated for writers, erasers, and readers using MEGA12 (Kumar et al., 2024). Model selection based on the Bayesian information criterion identified rtREV+G+F, rtREV+I+F, and WAG+G+I+F for the writer, eraser, and reader datasets, respectively. Node support was assessed using 2,000 bootstrap replicates.

### Taxonomic distribution of m^6^A-related gene copy numbers

Species names were mapped to NCBI taxonomy identifiers using ETE3 NCBITaxa, with manual correction of synonyms where required. NCBI lineage information was used to assign taxonomic ranks and organize the 119 species in a circular taxonomic display. For each species, copy numbers were calculated for five writer groups (MT-A70, Wtap, VIR_N, MTase-10, and RING-HC_HAKAI-like), one eraser group (ALKBH), and four reader groups (YTH, YTH+CPSF, CPSF, and RRM). Counts were derived from the locus-level representative protein sets prepared for the cross-species phylogenetic analyses. Within each group, one sequence was retained per species-specific gene identifier, prioritizing the longest isoform among proteins that passed the BLASTp and conserved-domain criteria. Each represented locus contributed one copy to its assigned group. The taxonomic grouping and ten group-specific copy-number heatmap tracks were visualized in iTOL. The taxonomic display does not represent a species phylogeny inferred from the m^6^A-related protein sequences.

### Gene structure and conserved domain architecture analysis

Gene structures and protein domain architectures of the identified *M. polymorpha* m^6^A-related genes were visualized using custom Python v3.13.9 scripts with pandas v2.3.3 and matplotlib v3.10.8. For gene-structure visualization, exon and CDS annotations were extracted from the MpTak_v7.1 GFF3 file using transcript parent IDs. Gene structures and domain architectures were visualized for 23 selected transcript and protein isoforms from 18 loci. The short MpECT12 isoform Mp3g00420.2, which lacks the YTH-domain region present in Mp3g00420.1, was excluded. The genomic start and end positions and strand of each selected mRNA feature were extracted from the GFF3 file to generate a transcript information table, and genomic coordinates were converted into transcript-oriented relative positions. Introns were defined as the intervals between adjacent exons, and UTRs were inferred as exon regions that did not overlap CDSs. All 23 selected transcripts had exon annotations, so the CDS-as-exon fallback was not used for this figure. In the final gene-structure plot, UTRs were shown as blue boxes, CDSs as yellow boxes, and introns as black lines. For domain visualization, the 23 selected protein sequences were extracted from the MpTak_v7.1 protein FASTA file and searched against a local Conserved Domain Database using RPS-BLAST with an E-value cutoff of 1 X 10^-3^. Hits to ten specified CDD identifiers were retained in a script-generated tab-delimited file. Protein lengths were calculated from the extracted protein FASTA file, and the RPS-BLAST query start and end positions were plotted relative to the full-length protein sequences. Overlapping hits from the two selected YTH domain models were collapsed when their overlap covered at least 80% of the shorter hit, prioritizing CDD identifier 410979. Domain names were simplified for plotting, while the original domain descriptions were retained in the tab-delimited RPS-BLAST results. Protein backbones were represented as grey horizontal bars, and annotated domains were displayed as colored rectangles according to domain type.

### Chromosomal location and collinearity analysis

The chromosomal locations of the 18 candidate m^6^A-related genes in *M. polymorpha* were determined from the MpTak_v7.1 GFF3 annotation. The corresponding genome assembly, v1.1 gene annotation, and primary-transcript protein dataset of *Marchantia quadrata* var. MQ were obtained from JGI Phytozome (Goodstein et al., 2012). For *M. polymorpha*, primary protein sequences were mapped to their parent gene loci, with one sequence retained per locus. For *M. quadrata*, the primary-transcript-only protein dataset was used. Transcript-level identifiers of the candidate *M. polymorpha* m^6^A-related genes were collapsed to their corresponding gene loci, and species-specific prefixes (MP_ and MQ_) were added to gene and chromosome identifiers to prevent identifier conflicts between the two genomes. Gene positions corresponding to the representative protein sequences from both genomes were extracted from the respective genome annotations and formatted as positional input for MCScanX v1.0.0.

Bidirectional protein similarity searches between the genome-wide representative protein sets of the two species were performed using DIAMOND v2.1.9 in BLASTP sensitive mode, with an E-value threshold of 1 X 10^-3^ and a maximum of 20 target sequences per query (Buchfink et al., 2021). The results from both search directions were combined, and interspecific collinear blocks were identified using MCScanX with the parameters -b 2 -s 3 -e 1e-5 -m 30 -a (Wang et al., 2012). These settings restricted the analysis to interspecific blocks, required a minimum of three anchor gene pairs per block, applied a block E-value threshold of 1 X 10^-5^, and set the maximum gap parameter to 30. The -a option limited output to the pairwise collinearity file. Duplicate *M. polymorpha*-*M. quadrata* collinear gene pairs were collapsed before downstream analysis.

Collinear pairs containing one of the 18 candidate *M. polymorpha* m^6^A-related genes were grouped according to whether the candidate belonged to the writer-related, ALKBH-family, or reader-related group. Pairs whose *M. polymorpha* member was not among the 18 candidates were classified as background relationships. The visualization included *M. polymorpha* chromosomes 1-8, U, and V and *M. quadrata* chromosomes 1-9. Collinear relationships were visualized using the Multiple Synteny Plot module of TBtools-II v2.498 (Chen et al., 2023). Candidate genes and their detected *M. quadrata* collinear counterparts were highlighted in red, green, and purple for the writer-related, ALKBH-family, and reader-related groups, respectively. Links involving these highlighted loci were displayed in the corresponding colors, including links connecting a highlighted *M. quadrata* locus to a non-candidate *M. polymorpha* gene. All other links were displayed in light grey. Only pairs containing one of the 18 candidate *M. polymorpha* genes were counted as candidate-associated collinear pairs. This analysis was used to assess conservation of chromosomal context and did not independently establish strict one-to-one orthology or the direction of gene duplication or loss.

### Identification of *cis*-regulatory elements in the putative promoter regions

For *cis*-regulatory element analysis, 2-kb genomic regions immediately upstream of the annotated 5′ end of each selected transcript model of the identified *M. polymorpha* m^6^A-related genes were extracted using the MpTak_v7.1 genome assembly and corresponding annotation. Upstream sequences were extracted according to transcript orientation, with sequences from negative-strand transcripts reverse-complemented. For loci represented by multiple transcript models in the curated dataset, a separate promoter entry was retained for each transcript model, resulting in 23 transcript-level promoter entries corresponding to 18 candidate m^6^A-related loci. Entries from the same locus could represent identical or overlapping upstream regions.

Putative *cis*-regulatory elements within the 2-kb promoter sequences were identified using PlantCARE (accessed 27 September, 2026) (Lescot et al., 2002; Rombauts et al., 1999). Site-level predictions were compiled from the PlantCARE output and processed using custom Python scripts. PlantCARE hits were retained for functional classification when their annotations could be assigned to one of the functional subcategories listed in Table 2. Hits outside these subcategories were excluded from the functional summaries. For Figure 5A, named motif hits were plotted regardless of whether they could be assigned to these subcategories; unnamed motif hits were omitted.

**Table 2.**
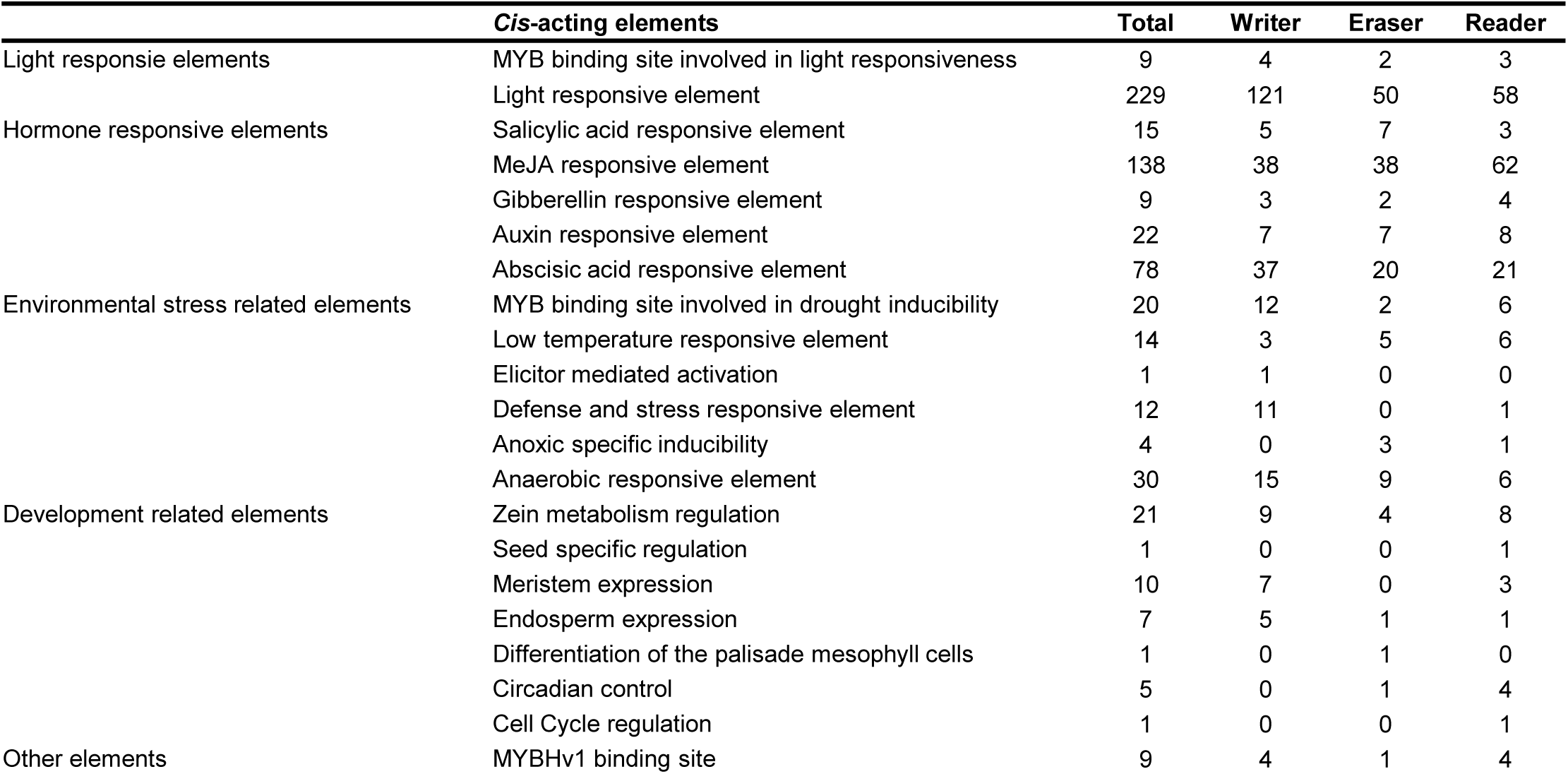
Predicted *cis*-elements in putative promoters of *M. polymorpha* candidate m^6^A-related genes. Predicted motif occurrences in the 2-kb upstream sequences of 23 transcript entries representing 18 loci are grouped by 21 PlantCARE functional subcategories. The Total column gives the number of predicted occurrences of each element across all entries; Writer, Eraser, and Reader give the corresponding counts within each candidate group. Across the listed elements, 636 occurrences were recorded: 282 in writer, 153 in eraser, and 201 in reader promoter entries. These are motif counts, not counts of genes with a motif. When transcript models have identical or overlapping upstream regions, their predictions contribute separately to the transcript-entry totals. MeJA, methyl jasmonate; MYB and MYBHv1, motif names used in the PlantCARE annotations. The functional categories are predictions and do not establish regulatory activity in *M. polymorpha*.

The retained motif hits were grouped into five major categories based on their PlantCARE functional annotations: light-responsive, hormone-responsive, environmental stress-related, development-related, and other elements (Table 2). MYBHv1-binding sites were assigned to the other-elements category. These annotations do not establish regulatory functions in *M. polymorpha*. To characterize motif distributions, the positions and identities of individual predicted elements were plotted along each 2-kb upstream sequence, from its distal 5′ end toward the annotated transcript 5′ end. A promoter-entry-by-functional-subcategory count matrix was generated by counting retained motif hits assigned to each functional subcategory within each transcript-level promoter entry and was visualized as a heatmap (Fugure 5B). Counts within each entry were additionally aggregated into the five major functional categories and displayed as stacked horizontal bar plots (Figure 5C). Totals reported in Table 2 were calculated across the 23 transcript-level promoter entries. All data processing and visualization were performed using custom Python scripts implemented in Python v3.13.9 with pandas v2.3.3, NumPy v2.3.5, matplotlib v3.10.8, and seaborn v0.13.2.

### Expression analysis across tissues and developmental stages

Expression data in transcripts per million (TPM) for the 18 candidate *M. polymorpha* m^6^A-related genes, comprising seven writer-related genes, five ALKBH-family genes, and six reader-related genes, were obtained from the MarpolBase Expression database (MBEX; accessed 9 September 2026) (Kawamura et al., 2022). The 18 candidate loci were matched to MBEX entries by their MarpolBase gene IDs, and gene-level TPM values were retrieved for expression analysis. The dataset included 16 tissue and developmental-stage conditions, covering a spore germination time course (0-96 h), vegetative and meristematic tissues, male and female reproductive tissues or cells, and early sporophyte tissue. Each tissue or developmental-stage condition was represented by two to four biological replicates. Replicate-level TPM values for the 18 candidate genes across the 16 tissue and developmental-stage conditions, together with their gene names and MarpolBase gene IDs, are provided in Supplementary Table S2.

Sample labels were standardized before analysis. Expression values from individual replicates were log_2_ (TPM + 1)-transformed and subsequently averaged within each tissue or developmental stage. No gene-wise Z-score transformation was applied. The heatmap display scale was set from -1 to 8, with values outside this range displayed at the corresponding limit. Separate heatmaps were generated for writers, erasers, and readers using the pheatmap and grid packages in R. Genes were hierarchically clustered within each functional class, whereas tissues and developmental stages were arranged in a predefined biological order without column clustering.

### RNA-seq data processing and differential expression analysis under abiotic stress

Raw RNA-seq data were obtained from ArrayExpress under accession number E-MTAB-11141 (Tan et al., 2023). The analyzed dataset comprised seven single-stress conditions (cold, heat, osmotic stress, salt stress, nitrogen deficiency, darkness, and high light), 18 pairwise stress combinations, and two unstressed control groups from independent experimental batches, designated Control D2 and Control H2. Sample accession numbers, treatment details, batch assignments, and biological replicate numbers are provided in [Table S X]. Zeitgeber time-course samples were excluded. Transcript abundance was quantified using Salmon in quasi-mapping mode against the M. polymorpha Tak-1 v5.1 cDNA reference transcriptome (Patro et al., 2017).

For transcript-to-gene mapping, only transcript identifiers ending in .1 were retained from the Tak-1 v5.1 cDNA annotation. Salmon abundance estimates were imported into R using tximport and summarized at the gene level using the transcript-to-gene mapping table, with countsFromAbundance = “lengthScaledTPM” (Soneson et al., 2015). Gene identifiers in the v5.1 reference were matched to the candidate loci identified using the v7.1 annotation.

Genes with fewer than 10 estimated counts summed across all analyzed samples were excluded before model fitting; all 18 candidate m^6^A-related genes passed this prefilter. A DESeq2 model was fitted to all retained genes across the 27 sample groups. Each stress condition was compared separately with Control D2 and Control H2 using Wald tests. Within each contrast, *P* values were adjusted using the Benjamini–Hochberg method before extracting the results for the 18 candidate genes (Love et al., 2014). Under each stress condition, a candidate gene was classified as consistently differentially expressed when both control comparisons yielded adjusted *P* values < 0.05 and log_2_ fold changes with the same sign. For each gene–stress combination meeting these criteria, the arithmetic mean of the log_2_ fold changes from the two control comparisons was used for heatmap visualization. Combinations that did not meet the criteria were displayed as zero, indicating failure to meet the selection criteria rather than an estimated absence of expression change. Separate heatmap panels were generated for single-stress and pairwise-stress conditions using pheatmap v[version] and grid in R v[version]. Both panels used the same asymmetric color scale ranging from -10 to +3, with white assigned to zero, red to positive values, and blue to negative values. Within each panel, genes were hierarchically clustered separately within the writer-related, ALKBH-family, and reader-related groups.

### qRT-PCR analysis

Quantitative reverse-transcription PCR (qRT-PCR) was performed to analyze transcript levels of the 18 candidate *M. polymorpha* m^6^A-related genes. Total RNA was extracted from thalli, gemmae, and gemma cups for tissue comparisons and from control and stress-treated plants for stress-response analyses using the Complex Plant RNA Extraction Kit (Adamas Life, G8028). cDNA was synthesized from 1 μg of total RNA using the RevertAid First Strand cDNA Synthesis Kit (Thermo Fisher Scientific, K1622). qRT-PCR was conducted using ChamQ Universal SYBR qPCR Master Mix (Vazyme, Q711-03) on a StepOnePlus Real-Time PCR system (Applied Biosystems). *MpSAND* was used as an internal reference gene (Saint-Marcoux et al., 2015). Relative transcript levels were calculated using the 2^-^ΔΔCt method, with *MpSAND* as the internal reference gene and thallus samples or the corresponding unstressed controls as calibrators for tissue comparisons or stress-response analyses, respectively (Livak & Schmittgen, 2001). Three independent biological replicates were analyzed for each condition, with three technical replicates performed for each biological replicate.

For stress-response analyses, plants were subjected to 40 mM NaCl or 100 mM mannitol treatment, cold treatment at 3 °C for 24 h, heat treatment at 33 °C for 24 h, or darkness for 3 days before sample collection. Primer sequences used for qRT-PCR are provided in Supplementary Table S3. Previously published primer sets for *MpSAND* and selected stress-responsive marker genes, including *MpFAD3*, *MpHSP17.8A1*, *MpLEA-like4*, *MpCIPK-B*, *MpCBL-A*, *MpABI3A*, *MpABI5B*, *MpSIG1*, and *MpSIG2*, were adopted from previous studies (Nishihama et al., 2016; Saint-Marcoux et al., 2015; Takemura et al., 2012; Tanaka et al., 2018; Tansley et al., 2023; Ueda et al., 2013), with the corresponding source references provided in Supplementary Table S3.

### Protein structure prediction and domain visualization

Three-dimensional structural models of 21 of the 23 candidate m^6^A-related protein isoforms were obtained from the AlphaFold Protein Structure Database (Varadi et al., 2022). Precomputed models from the Tak-1 proteome were used for all proteins except MpVIR-1 (Mp1g10290.1) and MpVIR-2 (Mp1g10290.2), for which models were unavailable. The structures of MpVIR-1 and MpVIR-2 were therefore predicted *de novo* using ColabFold, with multiple sequence alignments generated using MMseqs2 and five recycles used for model refinement. All other prediction parameters were maintained at their default settings (Mirdita et al., 2022; Steinegger & Soding, 2017).

Predicted structures were visualized in PyMOL v2.5.0 using cartoon representation. For visualization only, low-confidence residues were hidden according to the per-residue predicted local distance difference test (pLDDT) scores encoded in the B-factor column of each PDB file (Varadi et al., 2022). Protein-specific pLDDT thresholds ranging from 20 to 70 were applied to reduce the display of poorly resolved terminal, linker, and loop regions while retaining confidently folded regions. Conserved domains were identified using CD-Search against the NCBI Conserved Domain Database with RPS-BLAST, and the resulting domain boundaries were mapped onto the corresponding structural models (Camacho et al., 2009; Lu et al., 2020). MT-A70, Wtap, VIR_N, RING-HC_HAKAI-like, MTase-10, ALKBH, YTH, CPSF, and RRM domains were distinguished using a fixed color scheme, whereas regions without annotated domains were displayed in light grey.

All structures were rendered against a white background using consistent PyMOL settings. Individually rendered structures were assembled into a composite panel to compare the domain-associated structural features of writer, eraser, and reader proteins (Figure 10).

### Predicted protein-interaction network analysis

Predicted functional associations involving 18 representative candidate *M. polymorpha* m^6^A-related proteins, with one protein selected per locus, were retrieved from STRING v12.0 using the uploaded *M. polymorpha* proteome dataset (organism upload ID: STRG0A11VKF) (Szklarczyk et al., 2023). For loci with multiple annotated isoforms, the .1 isoform was used. Candidate proteins from the writer-related, ALKBH-family, and reader-related groups were submitted as query nodes. The minimum required interaction score was set to 0.4, and the maximum numbers of additional interactors in the first and second shells were set to 20 and 5, respectively. All other STRING parameters were maintained at their default settings.

The resulting network was visualized using Cytoscape v3.10.4, and unconnected query proteins were retained to show candidate proteins for which no association meeting the selected threshold was identified (Shannon et al., 2003). Network edges were interpreted as predicted functional protein associations supported by the evidence sources integrated in STRING and not necessarily as direct physical protein-protein interactions (Szklarczyk et al., 2023).

## Statistical analysis

Statistical analyses were performed using GraphPad Prism 10.5.0. For qRT-PCR comparisons among tissues and among control and abiotic-stress conditions, data were analyzed by ordinary one-way analysis of variance (ANOVA), assuming equal variances, followed by Tukey’s multiple-comparisons test for all pairwise comparisons. For two-group comparisons between the control and an individual stress treatment in the stress-marker validation experiment, a two-tailed unpaired Welch’s t-test was used. For Tukey’s test, the family-wise significance level was set at α = 0.05, and multiplicity-adjusted *P* values were reported. Significance was denoted as ns (*P* ≥ 0.05), * (*P* < 0.05), ** (*P* < 0.01), *** (*P* < 0.001), and **** (*P* < 0.0001).

## Results

### Genome-wide analysis reveals conserved plant m^6^A-related gene families and identifies 18 genes in *Marchantia polymorpha*

To define the evolutionary landscape of plant m^6^A regulation, we first performed a genome-wide identification of putative writer, eraser, and reader proteins across 119 plant genomes. Candidate genes were identified using *Arabidopsis thaliana* m^6^A-related proteins as reference queries and retained when their encoded proteins contained conserved domains consistent with their family assignments. The annotated domains included MT-A70, Wtap, VIR_N, Methyltransf_10 (MTase-10), RING-HC_HAKAI-like, 2OG-FeII_Oxy_2 (ALKBH), YT521-B homology (YTH), cleavage and polyadenylation specificity factor (CPSF), and RRM_ist3_like (RRM) domains. This survey showed that the three functional classes of m^6^A-related proteins were broadly represented across the sampled plant lineages, although individual families differed substantially in copy number among species (Figure 2). Canonical writer components generally occurred at relatively low copy numbers, whereas ALKBH-family erasers and reader proteins showed broader copy-number variation. These patterns are consistent with broad conservation of plant m^6^A regulatory components and lineage-dependent variation in gene family size (Zhang et al., 2024). Together, the copy-number survey provides a broad evolutionary context for defining the m^6^A-related gene repertoire in *M. polymorpha*.

**Figure 2.**
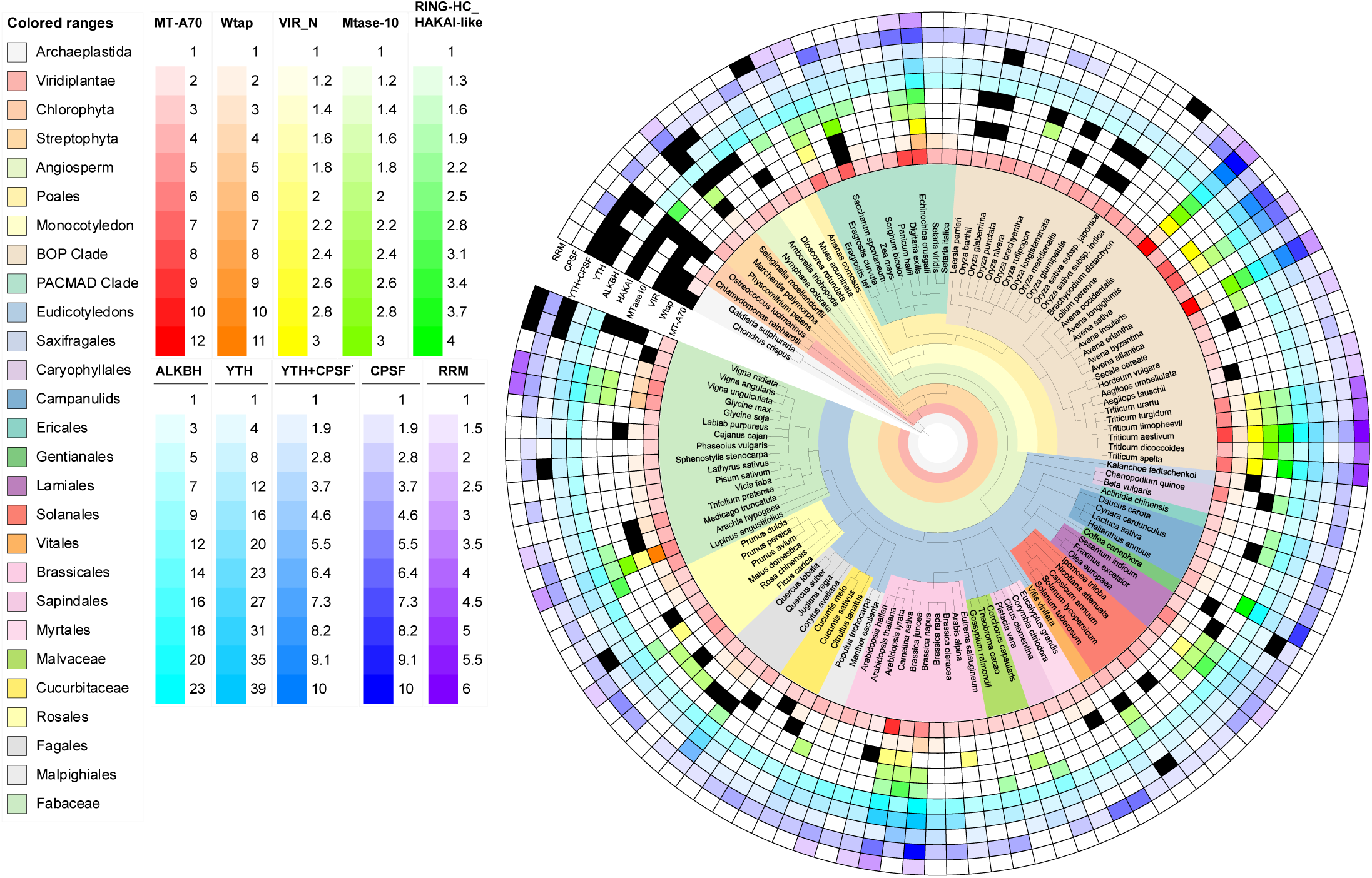
Distribution of candidate m^6^A regulator families across 119 plant species. Each radial column represents a species. From the outermost to innermost ring, the ten rings show copy numbers for RRM_ist3_like (RRM), Cleavage and polyadenylation specificity factor (CPSF), YTH plus CPSF (YTH+CPSF), YTH, 2OG-FeII_Oxy_2 (ALKBH), RING-HC_HAKAI-like, Methyltransf_10 (MTase-10), VIR_N, Wtap, and MT-A70 categories. Within each category, stronger color intensity indicates a higher value according to its separate gradient in the key; black indicates zero detected copies. Colored taxonomic sectors correspond to the lineages identified in the left-hand key, and the central tree depicts relationships among the sampled species. BOP and PACMAD designate the Bambusoideae-Oryzoideae-Pooideae and Panicoideae-Aristidoideae-Chloridoideae-Micrairoideae-Arundinoideae-Danthonioideae grass clades, respectively. One protein isoform per locus was retained for cross-species counts. The central tree and colored sectors represent NCBI taxonomy and do not constitute an m^6^A-protein phylogeny.

Maximum-likelihood phylogenetic analyses were performed for the writer, eraser, and reader datasets (Supplementary Figures 1-3). Writer proteins separated into MTA-, MTB-, MTC-, FIP37-, VIR-, FIO1-, and HAKAI-related clades, whereas eraser proteins clustered within ALKBH-related clades. The reader dataset included proteins annotated in four domain categories: YTH, YTH+CPSF, CPSF, and RRM. *M. polymorpha* candidates were placed within the corresponding plant clades, supporting their family assignments.

Using this identification and domain-validation strategy, we identified 18 candidate m^6^A regulatory genes in *M. polymorpha*, including seven writers, five erasers, and six readers (Table 1). The writer genes comprised *MpMTA*, *MpMTB*, *MpMTC*, *MpFIP37*, *MpVIR*, *MpFIO1*, and *MpHAKAI*. The eraser genes comprised *MpALKBH1A*, *MpALKBH1D*, *MpALKBH6*, *MpALKBH8B*, and *MpALKBH9C*. The reader genes comprised *MpECT2*, *MpECT12*, *MpCPSF30-L1*, *MpCPSF30-L3*, *MpCPSF30-L4*, and *MpCPSF30-L5*. Five loci—*MpMTA*, *MpFIP37*, *MpVIR*, *MpFIO1*, and *MpALKBH1D*—were each represented by two annotated transcript models, yielding 23 transcript models across the 18 loci. Nuclear localization was predicted for many of the identified proteins. Cytoplasmic or organellar localization was also predicted for several proteins, including MpFIO1, MpALKBH6, MpALKBH8B, MpALKBH9C, MpCPSF30-L4, and MpCPSF30-L5, by one or more tools (Table 1). These predictions suggest potential differences in subcellular distribution among the proteins. This set of 18 *M. polymorpha* m^6^A-related genes provided the basis for subsequent analyses of gene structure, domain architecture, chromosomal distribution and collinearity, putative promoter elements, expression patterns, predicted protein structures, and predicted protein associations.

### Gene structure and domain analyses support the annotation of *M. polymorpha* candidate m^6^A regulators

We next examined the exon–intron structures and conserved domain composition of the identified *M. polymorpha* candidate m^6^A regulators. Exon–intron structures and protein domain architectures are shown in Figure 3A and B, respectively. Exon–intron organization varied among the three functional classes, although several candidates within related phylogenetic groups showed comparable structural features (Figure 3A). The writer genes displayed substantial structural diversity. Among the writer candidates, MpVIR encoded one of the largest predicted proteins and showed an extended gene model with numerous exons, whereas MpMTC and MpHAKAI displayed more compact gene structures. MpFIP37 and MpFIO1 were each represented by two annotated transcript models, with similar overall exon–intron organization between the two models at each locus. Among the reader proteins, MpCPSF30-L3 had the greatest predicted length (Figure 3B). Overall, these genes showed diverse exon–intron structures, with similarities among some phylogenetically related members.

**Figure 3.**
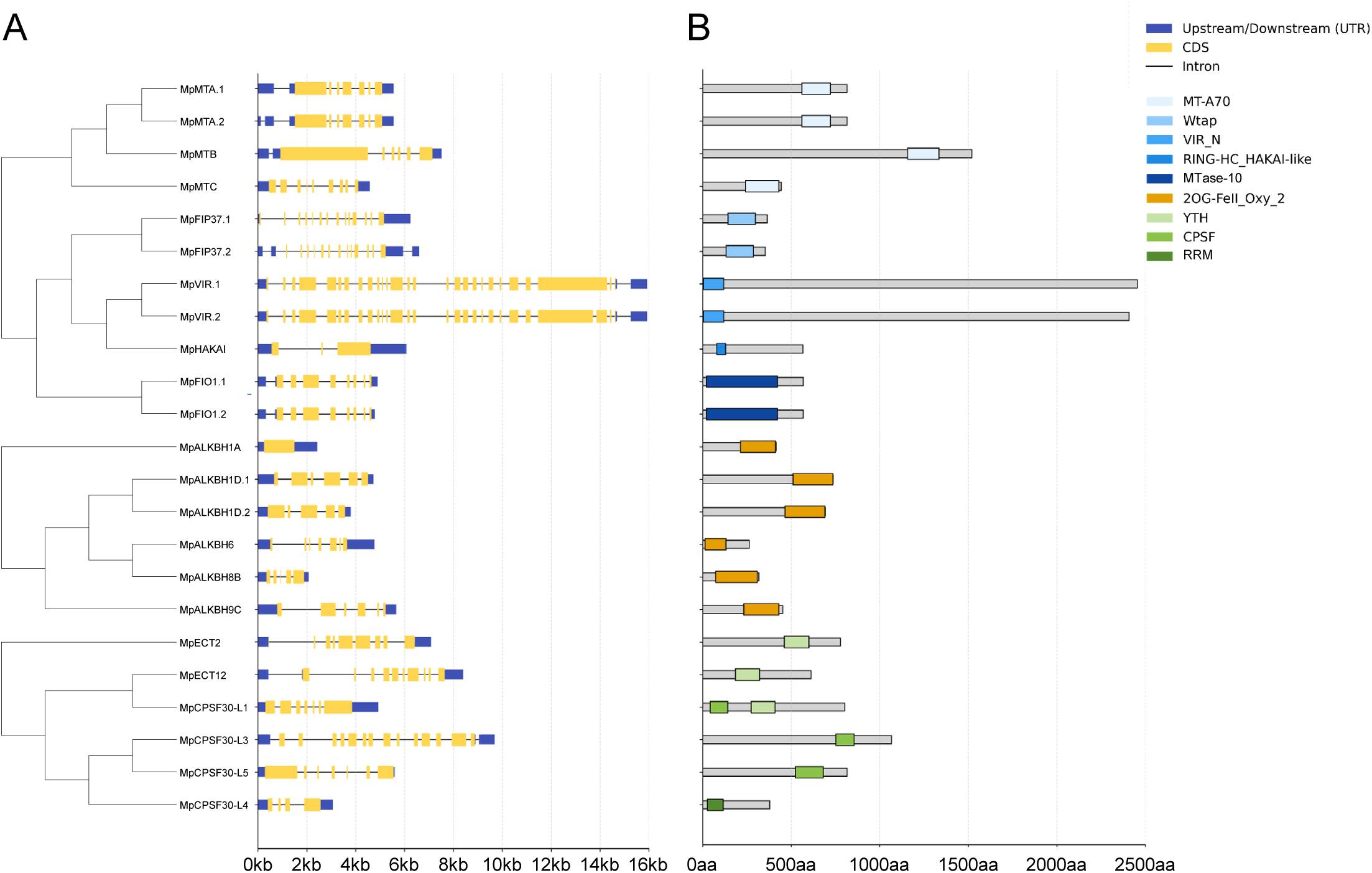
Exon-intron structures and conserved protein domains of *M. polymorpha* candidate m^6^A regulators. **a** Separate maximum-likelihood gene trees and transcript-specific gene structures of the writer, eraser, and reader candidates. Each row represents a selected transcript isoform. Yellow boxes indicate coding sequences (CDS), dark-blue boxes untranslated regions (UTRs), and thin black lines introns. The horizontal scale is in kilobases (kb). **b** Conserved domains mapped onto the proteins encoded by the transcript models in (A). Grey bars indicate full protein lengths. Colored blocks indicate MT-A70 (very pale blue), Wtap (pale blue), VIR_N (blue), RING-HC_HAKAI-like (medium blue), MTase-10 (dark blue), 2OG-FeII_Oxy_2 (orange), YTH (yellow-green), CPSF (green), and RRM (dark green) domains, as keyed in the panel. The lower scale is in amino acids (aa).

Conserved-domain analysis revealed the domain architectures of the identified proteins (Figure 3B). Among the writer proteins, MpMTA, MpMTB, and MpMTC contained MT-A70 domains, whereas MpFIP37, MpVIR, MpFIO1, and MpHAKAI contained Wtap, VIR_N, MTase-10, and RING-HC_HAKAI-like domains, respectively. All eraser proteins contained ALKBH-related domains. Proteins assigned to the reader group contained YTH, CPSF, and/or RRM domains. Together, the domain annotations and phylogenetic analyses supported the family assignments of the 18 identified m^6^A-related genes. We next examined the chromosomal distribution of these genes and their collinear relationships with loci in *Marchantia quadrata*.

### Chromosomal distribution and collinearity analysis reveal conserved genomic context for eight *M. polymorpha* m^6^A-related genes

To place the *M. polymorpha* candidates in a genomic and genus-level comparative context, we examined their chromosomal distribution and interspecific collinearity with the closely related species *M. quadrata*. The 18 candidate m^6^A-related genes, comprising seven writers, five erasers, and six readers, were distributed across multiple *M. polymorpha* chromosomes rather than being concentrated within a single genomic region (Figure 4). Specifically, the candidates were located on chromosomes 1, 3, 4, 6, 7, and 8, whereas no candidate m^6^A-related genes were identified on chromosomes 2, 5, U, or V. Chromosome 1 contained the largest number of candidates, with eight loci comprising five writers, two erasers, and one reader. Chromosome 3 contained three candidates. Chromosome 4 contained one candidate, whereas chromosomes 6, 7, and 8 each contained two.

**Figure 4.**
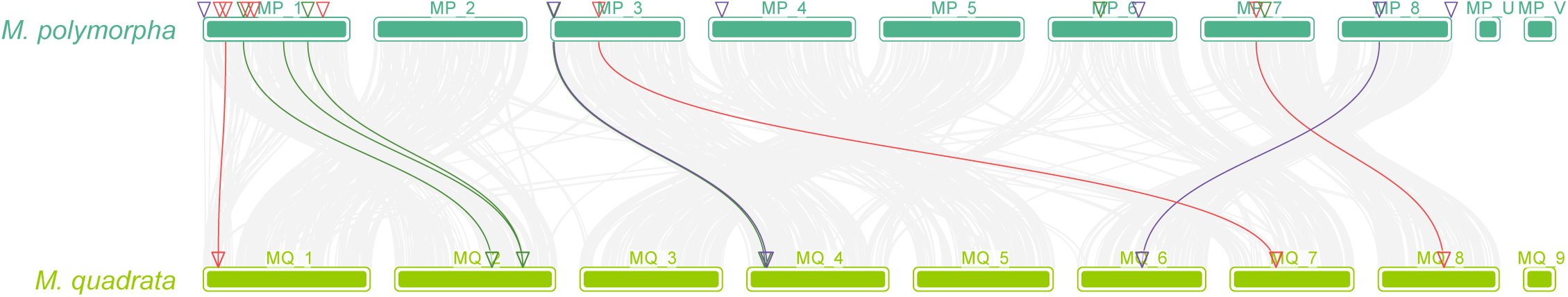
Chromosomal positions and interspecific collinearity of candidate m^6^A-related genes in *Marchantia*. Chromosomes of *M. polymorpha* (MP_, teal bars, top) and *Marchantia quadrata* (MQ_, lime-green bars, bottom) are shown in reference-assembly order. The input data input contains 5,560 interspecific collinear gene pairs. Light-grey curves represent 5,551 pairs without highlighted endpoints. Eight colored curves connect an *M. polymorpha* candidate to an *M. quadrata* locus: three writer-associated links in red, three eraser-associated links in green, and two reader-associated links in purple. A ninth colored curve, also green, connects *M. polymorpha* Mp1g16590 to *M. quadrata* Maqua.2G183700. Mp1g16590 is not among the 18 candidates in Table 1; this curve is highlighted because its *M. quadrata* endpoint is also linked to the eraser candidate Mp1g21480. Correspondingly colored triangles mark candidate loci and their detected collinear counterparts. MP_ and MQ_ are species-specific chromosome prefixes; U and V designate the sex chromosomes labeled in the *M. polymorpha* assembly.

Genome-wide comparison between *M. polymorpha* and *M. quadrata* identified 5,567 interspecific collinear gene pairs. Among the 18 *M. polymorpha* m^6^A-related loci, eight were associated with detectable collinear relationships in *M. quadrata*, comprising three writer, three eraser, and two reader genes (Figure 4). Thus, detectable collinearity was retained for 3 of 7 writer loci, 3 of 5 eraser loci, and 2 of 6 reader loci under the criteria used in this analysis.

At the individual-gene level, the writer genes Mp1g04450, Mp3g09940, and Mp7g09040 formed collinear pairs with *M. quadrata* loci Maqua.1G029900, Maqua.7G060800, and Maqua.8G103000, respectively. Among the eraser candidates, Mp1g08100, Mp1g21480, and Mp3g00190 formed collinear pairs with Maqua.2G120300, Maqua.2G183700, and Maqua.4G032100, respectively. Two reader genes, Mp3g00420 and Mp8g06500, formed collinear pairs with Maqua.4G034600 and Maqua.6G082600, respectively. These relationships involved eight distinct loci in each species and were distributed across several chromosomes rather than being restricted to a single conserved chromosomal segment.

By contrast, no collinear counterpart was detected under the applied parameters for the remaining four writer genes (Mp1g03170, Mp1g08870, Mp1g10290, and Mp1g24270), two eraser genes (Mp6g08620 and Mp7g10560), or four reader genes (Mp1g00140, Mp4g02260, Mp6g15200, and Mp8g18820). The absence of a detected collinear counterpart under the applied criteria does not establish the absence of a homologous gene in *M. quadrata*. Together, these results identified eight M. polymorpha m^6^A-related loci with conserved genomic context in *M. quadrata*, complementing the sequence-based phylogenetic analyses.

### Putative promoters of *M. polymorpha* m^6^A-related genes contain abundant light-, hormone-, development-, and stress-associated *cis*-elements

To explore potential transcriptional regulation of *M. polymorpha* m^6^A-related genes during development and environmental responses, we analyzed predicted cis-regulatory elements in 2-kb genomic regions immediately upstream of the annotated 5′ ends of the transcript models of 23 transcript models representing 18 loci. These regions were treated as putative promoters, and the positions of predicted *cis*-elements were mapped for each transcript model (Figure 5A).

**Figure 5.**
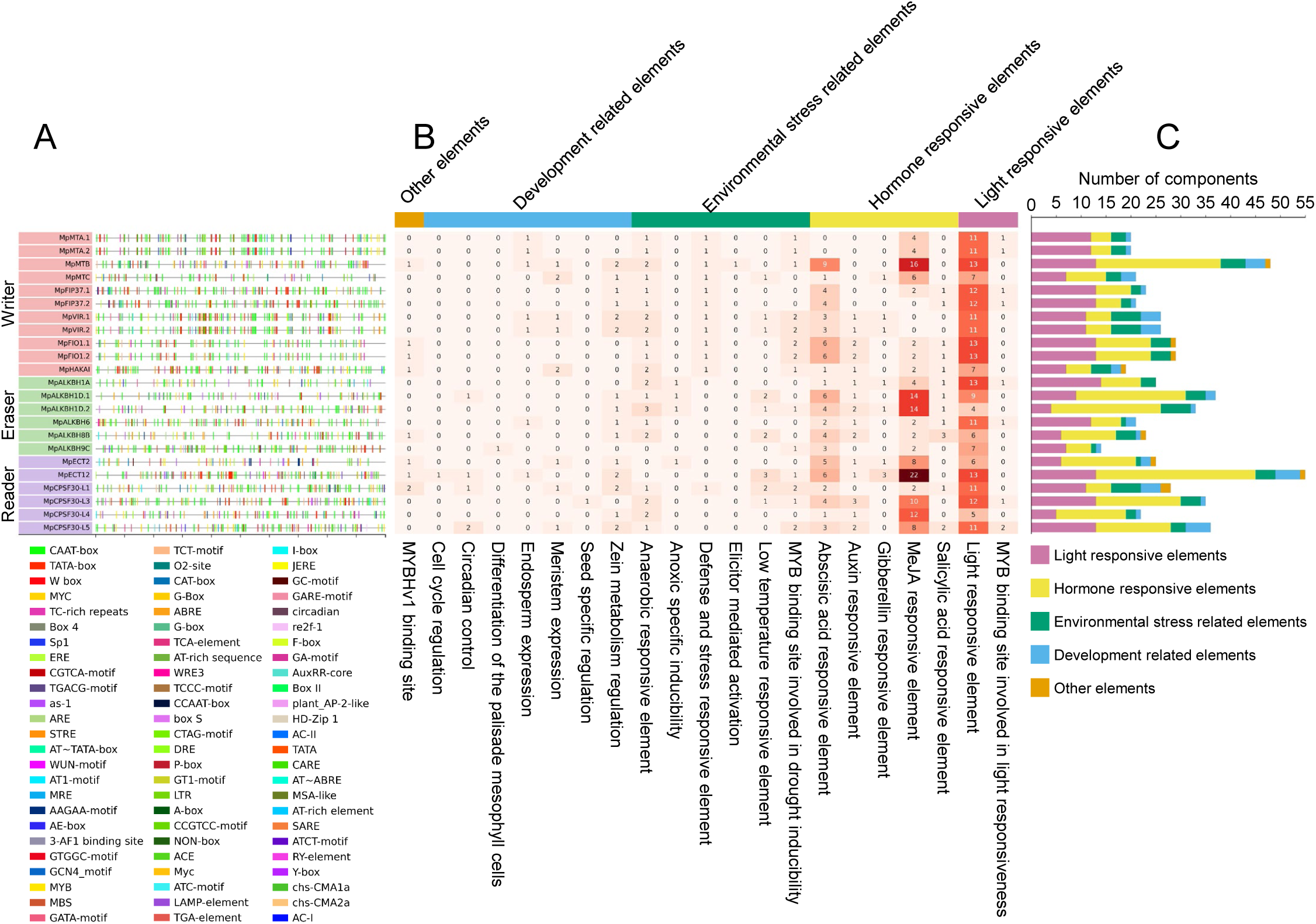
Predicted *cis*-elements in putative promoters of *M. polymorpha* candidate m^6^A-related genes. **a** Positions of 2,619 named motif occurrences predicted by PlantCARE within 2-kb upstream sequences of 23 selected transcript models representing 18 candidate loci. Each colored tick denotes a predicted motif; its color corresponds to the motif named in the key beneath the plot. The horizontal axis gives position in base pairs (bp) from the distal 5′ end toward the annotated transcript 5′ end. Separate rows for isoforms of the same locus represent the analyzed transcript-level promoter entries. Colored label backgrounds distinguish writer (pink), eraser (green), and reader (purple) entries. **b** Heatmap of 636 functionally classified motif occurrences across 21 annotated functional subcategories for the same 23 promoter entries. Darker red indicates more annotated occurrences, whereas pale or white cells indicate few or none. The colored strip above the heatmap identifies the five major categories; values in parentheses give their totals across 23 entries. **c** Stacked bars aggregate sites into light-responsive (purple), hormone-responsive (yellow), environmental stress-related (green), development-related (light blue), and other (orange) categories. The bar-plot axis gives the number of predicted motif occurrences. These are PlantCARE motif annotations, rather than experimentally established functions in *M. polymorpha*. MeJA, methyl jasmonate; ABA, abscisic acid; MYB, MYB-family binding motif.

In total, 636 predicted motif occurrences were retained for functional classification across the 23 putative promoter sequences, including 282 in writer-gene promoters, 153 in eraser-gene promoters, and 201 in reader-gene promoters (Table 2). These elements were grouped into five functional categories based on PlantCARE annotations: light-responsive, hormone-responsive, environmental stress-related, development-related, and other elements (Figure 5B). This classification provided an overview of the potential regulatory inputs associated with the identified m^6^A-related genes.

Hormone-responsive elements represented the most abundant major category, whereas the subcategory annotated as light-responsive element had the highest motif count. This subcategory accounted for 262 predicted motif occurrences, including 90 in writer-gene promoters, 74 in eraser-gene promoters, and 98 in reader-gene promoters. The motif class annotated as a light-responsive element was detected 238 times, including 125 sites in writer-gene promoters, 52 sites in eraser-gene promoters, and 61 sites in reader-gene promoters. Hormone-associated elements were also highly represented across the putative promoter regions. Methyl jasmonate (MeJA)-responsive and abscisic acid (ABA)-responsive elements accounted for 138 and 78 predicted motif occurrences, respectively. Auxin-responsive elements were also present, with 22 predicted sites across the analyzed promoter regions. Stress-associated motifs were detected across the putative promoters, including MYB binding sites associated with drought inducibility, low-temperature-responsive elements, defense/stress-responsive elements, anoxic inducibility elements, and anaerobic-responsive elements. Development-related and growth-associated motifs, including motifs annotated as zein metabolism-related motifs, meristem expression elements, endosperm expression elements, circadian elements, and cell cycle regulation elements, were also detected. Because several motif names are derived from annotations in seed plants, these elements should be interpreted as sequence motifs associated with regulatory categories rather than as direct evidence for the same biological processes in *M. polymorpha*.

The distribution of predicted *cis*-elements differed among individual m^6^A-related transcript-level promoter entries (Figures 5B and 5C). MpECT12 had the highest number of functionally classified occurrences, including 32 hormone-responsive annotations. Together, these motif annotations generate hypotheses about light-, hormone-, stress-, and development-associated regulations that require experimental testing. These predictions provide a basis for comparing promoter architecture with tissue- and stress-dependent expression patterns in the following sections.

### *M. polymorpha* m^6^A-related genes show tissue- and developmental-stage-dependent expression

We next examined the expression patterns of the 18 *M. polymorpha* m^6^A-related genes across 16 tissue and developmental-stage conditions using data from the MarpolBase Expression database (MBEX) (Kawamura et al., 2022). Heatmap analysis showed that all 18 genes had nonzero TPM values in at least one sampled condition, but their expression levels varied across samples (Figure 6). Several genes showed relatively broad expression across samples from the spore germination time course, thallus, reproductive tissues or cells, and early sporophytes. For example, *MpECT2* showed relatively high expression across the sampled conditions, whereas *MpCPSF30-L3* showed lower expression in the 9-day apical notch and higher expression in the antheridium. Others showed more restricted or condition-enriched expression patterns. Hierarchical clustering within the writer, eraser, and reader groups organized genes according to similarities in their expression profiles across the sampled tissues and developmental stages. These heatmap-based expression profiles provided context for the subsequent qRT-PCR analysis of tissue-associated expression. To further examine tissue-associated expression, we measured transcript levels in thallus, gemmae, and gemma cups using qRT-PCR (Figure 7). Most writer genes showed clear tissue-dependent expression patterns. *MpMTA*, *MpMTC*, *MpFIP37*, *MpVIR*, and *MpFIO1* transcript levels were higher in thallus than in gemmae or gemma cups. *MpMTB* showed lower expression in gemma than in thallus or gemmae cup, whereas *MpHAKAI* showed no significant difference among tissues. Among erasers, *MpALKBH1A* showed no significant difference among tissues, whereas *MpALKBH1D*, *MpALKBH6*, *MpALKBH8B*, and *MpALKBH9C* were more highly expressed in thallus than in gemmae or gemma cups. Reader genes also showed tissue-dependent expression: all six reader candidates were more highly expressed in thallus than in gemmae or gemma cups. *MpECT12* was more highly expressed in gemma cups than in gemmae, whereas *MpCPSF30-L3* was more highly expressed in gemmae than in gemma cups. Thus, the qRT-PCR results complemented the MBEX expression profiles by revealing gene-specific expression differences among the three tested tissues, with many writer, eraser, and reader genes showing higher expression in the thallus. Together, these expression patterns suggest potential roles for m^6^A-related genes in distinct vegetative and reproductive contexts in *M. polymorpha*.

**Figure 6.**
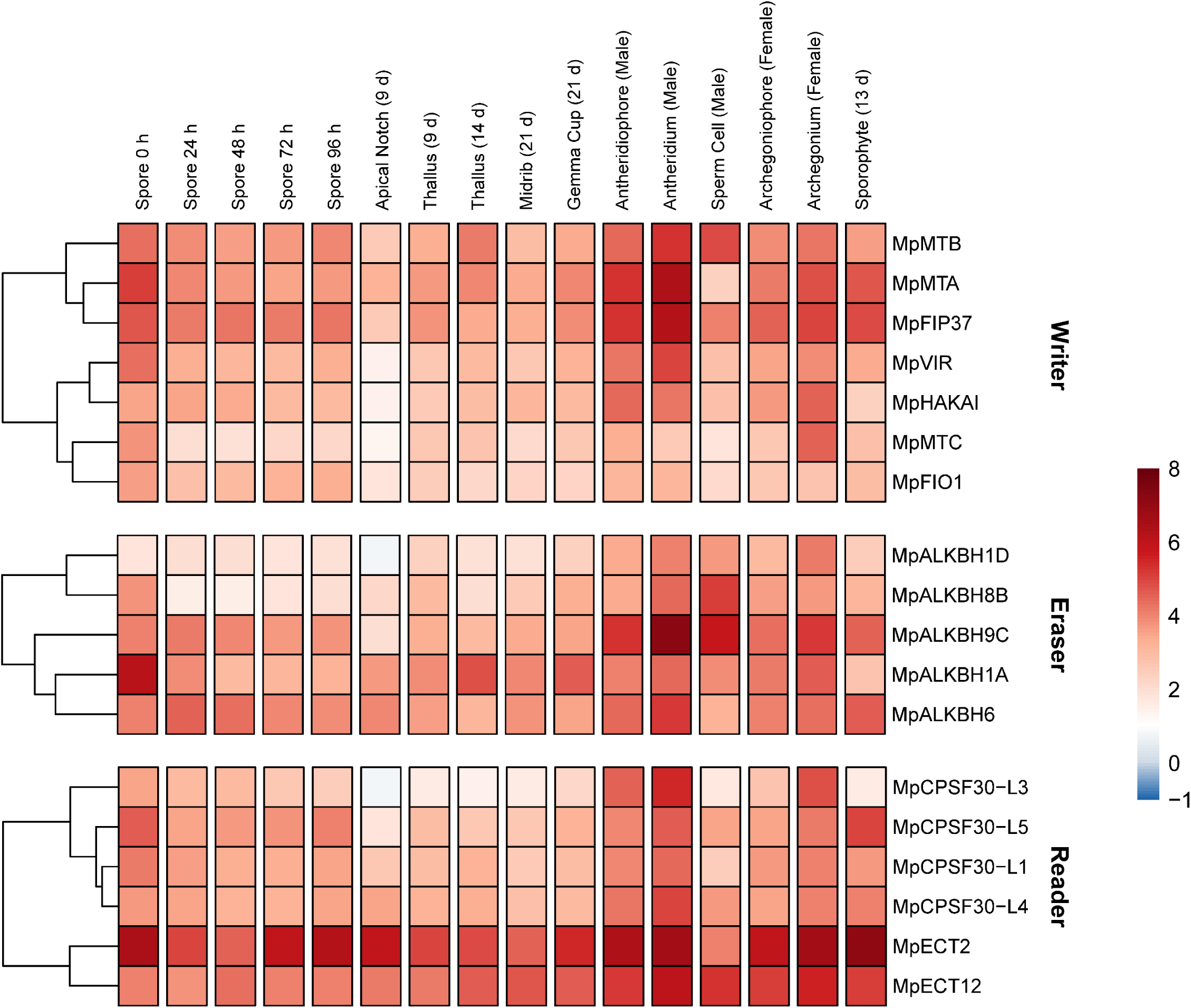
Expression of *M. polymorpha* candidate m^6^A-related genes across tissues and developmental stages. Writer, eraser, and reader genes are shown in separately clustered blocks, with dendrograms at left. Columns show the spore time course (0-96 h), apical notch, thallus, midrib, gemma cup, male and female reproductive tissues or cells, and sporophyte; sample identity, age, and sex are indicated above the heatmap. Values are condition means of replicate-level log_2_ (TPM + 1) values from the MarpolBase Expression database (MBEX) data; the display scale is bounded at -1 and 8. Red indicates higher transformed expression, blue lower expression, and white values near log_2_ (TPM + 1) = 1. h, hours; d, days.

**Figure 7.**
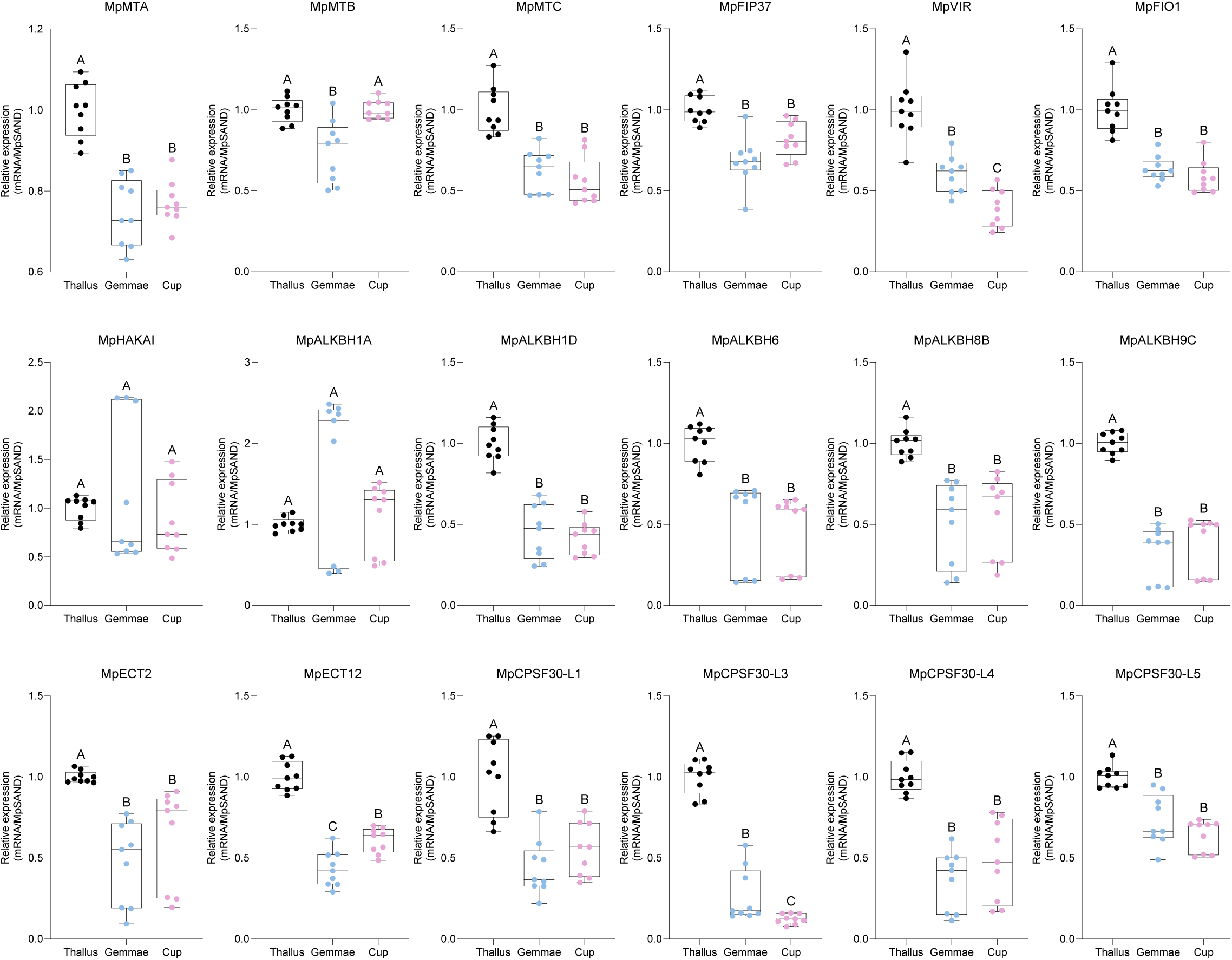
Tissue-associated expression of 18 *M. polymorpha* candidate m^6^A-related genes measured by qRT-PCR. Individual panels for the seven writer, five eraser, and six reader candidates are arranged in that order from left to right and top to bottom. Thallus is represented by black points, gemmae by blue points, and gemma cups - labeled Cup in the plots - by magenta points. *MpSAND* was used as the internal reference. Boxes span the first to third quartiles, center lines indicate medians, whiskers indicate the plotted ranges, and points show individual measurements. Statistical comparisons used ordinary one-way ANOVA followed by Tukey’s multiple-comparisons test for all tissue pairs. Groups without a shared uppercase letter differ at adjusted *P* < 0.05.

### Abiotic stress alters the expression of *M. polymorpha* m^6^A-related genes

We examined stress-associated expression changes in the 18 *M. polymorpha* m^6^A-related genes by reanalyzing published RNA-seq data (Tan et al., 2023). Under the seven single-stress conditions, writer, eraser, and reader genes displayed stress-dependent expression changes relative to unstressed controls (Figure 8A). The response was not uniform across the writer, eraser, and reader gene groups. Some writer genes showed increased expression under specific stresses, whereas others showed reduced expression. Eraser genes also showed divergent responses, suggesting that stress may differentially affect the expression of genes associated with m^6^A methylation and demethylation pathways. Reader genes also showed gene-specific increases and decreases across the tested stresses. Heatmaps for the 18 pairwise stress combinations showed expression patterns that differed from those observed under the corresponding individual stresses (Figure 8A, B). These data suggest that *M. polymorpha* m^6^A-related genes are associated with stress-responsive transcriptional programs. These RNA-seq analyses provided a broader context for the subsequent qRT-PCR analysis under selected abiotic stress conditions.

**Figure 8.**
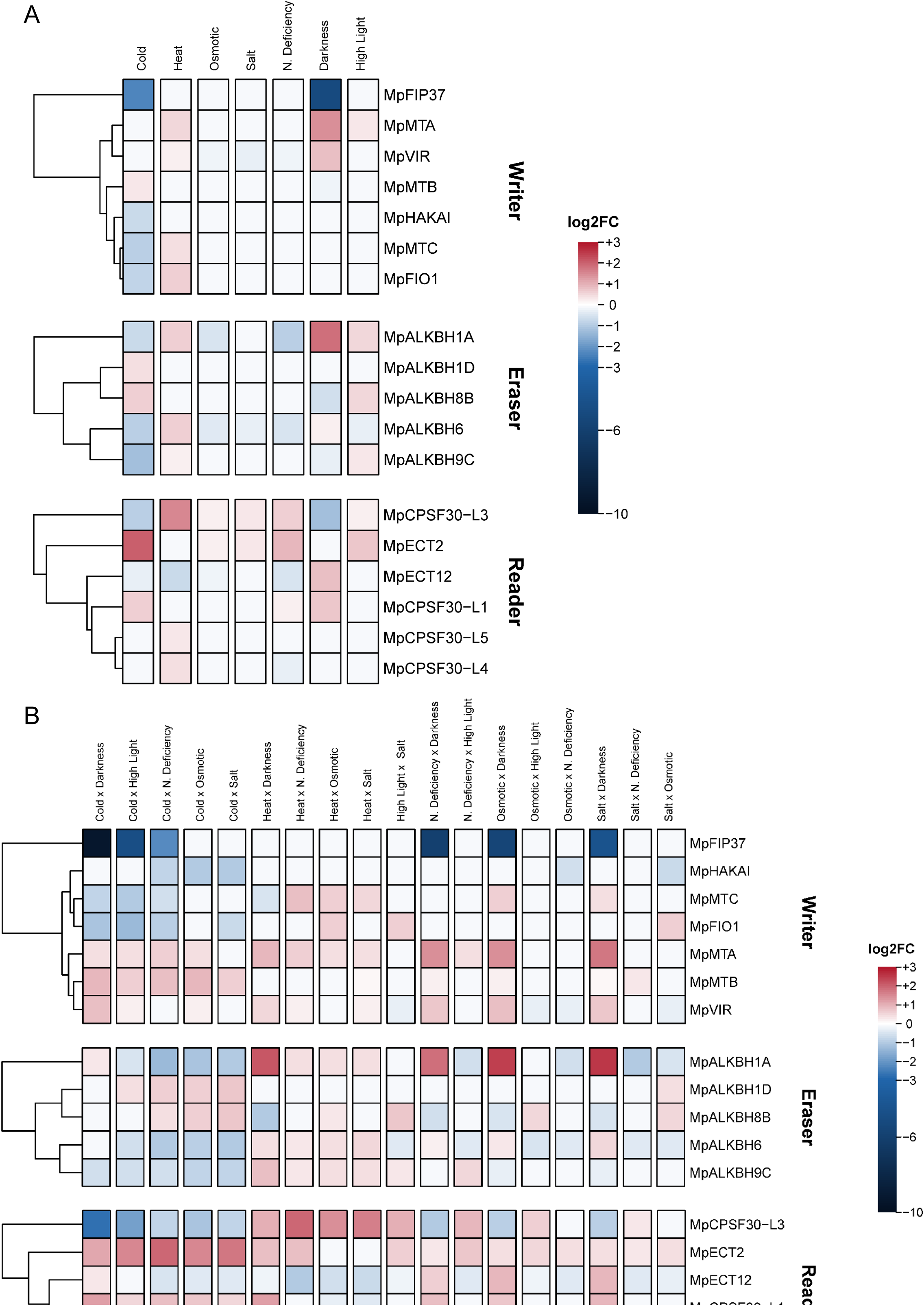
Differential expression of *M. polymorpha* candidate m^6^A-related genes under abiotic stress in RNA-seq data. **a** Single-stress conditions. **b** Pairwise stress combinations; “x” in a column label denotes a combined treatment. Writers, erasers, and readers are clustered separately. Cells show the mean log_2_ fold change (log_2_FC) relative to Control D2 and Control H2 when adjusted *P* < 0.05 and the direction of change agreed in both comparisons; otherwise, the displayed value is zero. Red denotes increased expression, blue decreased expression, and white a display value of zero, which does not necessarily indicate an estimated log_2_FC of zero. The color scale ranges asymmetrically from -10 to +3. N. deficiency, nitrogen deficiency; high light, elevated-light treatment.

Expression changes in stress-responsive marker genes confirmed responses to the selected treatments (Supplementary Figure 4). qRT-PCR analysis showed that many *M. polymorpha* m^6^A-related genes exhibited altered transcript levels relative to unstressed controls following exposure to cold, heat, osmotic stress, salt stress, or darkness (Figure 9). Darkness increased the transcript levels of multiple genes, including *MpMTA*, *MpMTC*, *MpVIR*, *MpFIO1*, *MpALKBH1A*, *MpECT2*, *MpCPSF30-L1*, and *MpCPSF30-L3*. Heat and osmotic stress also increased the expression of subsets of genes. For example, expression increased for *MpALKBH9C* under heat, for *MpALKBH6* under heat and osmotic stress, and for *MpECT12* under osmotic stress. Salt stress altered the expression of a subset of eraser and reader genes, including *MpALKBH1A*, *MpALKBH6*, *MpECT12*, and *MpCPSF30-L1*. By contrast, several genes showed reduced expression under cold stress, including *MpFIP37*, *MpMTA*, *MpMTC*, and *MpALKBH8B*. *MpHAKAI* showed a contrasting pattern relative to several other writer genes, with higher expression under normal or cold conditions than under salt or darkness. Together, the RNA-seq and qRT-PCR analyses revealed gene- and stress-dependent expression responses among *M. polymorpha* m^6^A-related genes.

**Figure 9.**
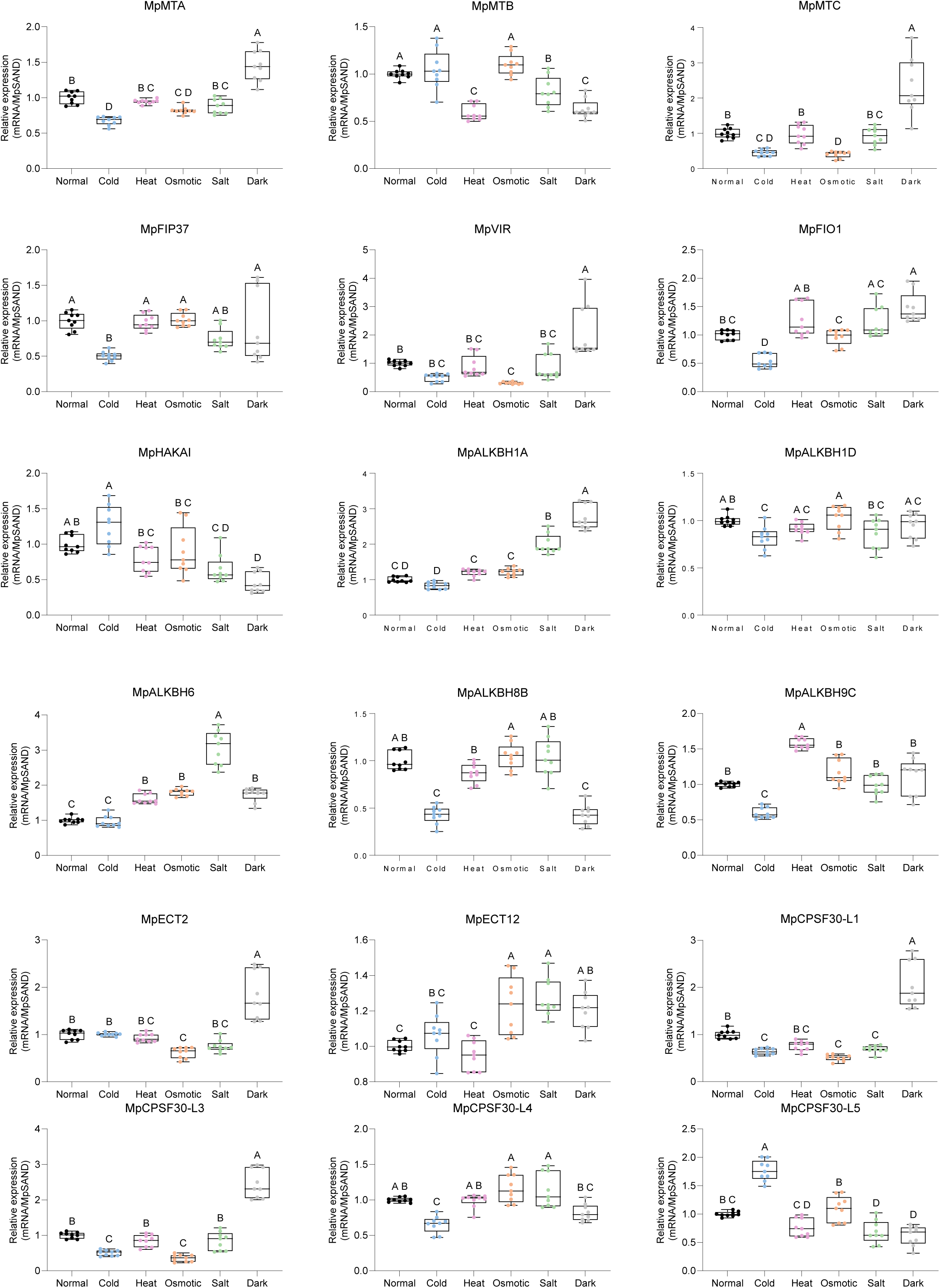
Expression of 18 *M. polymorpha* candidate m^6^A-related genes under abiotic stress measured by qRT-PCR. Treatments are Normal (black points), cold at 3 °C for 24 h (blue), heat at 33 °C for 24 h (pink), osmotic stress induced by 100 mM mannitol (orange), salt stress induced by 40 mM NaCl (green), and darkness for 3 d (grey). Expression is relative to the *MpSAND* internal reference. Boxes span the first to third quartiles, center lines indicate medians, whiskers indicate the plotted ranges, and points show individual measurements. Statistical comparisons used ordinary one-way ANOVA followed by Tukey’s multiple-comparisons test for all treatment pairs. Groups without a shared uppercase letter differ at adjusted *P* < 0.05.

### Predicted protein structures are consistent with conserved domain architectures in *M. polymorpha* m^6^A-related proteins

To visualize the structural context of the sequence-based domain annotations, we examined predicted protein models corresponding to the 23 annotated transcript models of the 18 *M. polymorpha* m^6^A-related loci. Twenty-one models were retrieved from the AlphaFold Protein Structure Database, and the two MpVIR isoforms were modeled using ColabFold (Figure 10). For each of the five loci with two annotated transcript models—MpMTA, MpFIP37, MpVIR, MpFIO1, and MpALKBH1D—the corresponding protein models are displayed separately under a shared gene label. Annotated domains were mapped onto the predicted models and highlighted using a fixed color scheme, whereas displayed residues outside these domains were shown in grey. Residues below the protein-specific pLDDT thresholds were omitted from the visualization. The mapped regions included MT-A70 domains in MpMTA, MpMTB, and MpMTC; Wtap, VIR_N, MTase-10, and RING-HC_HAKAI-like domains in MpFIP37, MpVIR, MpFIO1, and MpHAKAI, respectively; ALKBH-related domains in the eraser candidates; and YTH, CPSF, or RRM-related domains in the reader candidates (Figure 10). These predicted models provide a structural context for the sequence-based domain annotations and complement the phylogenetic and conserved-domain analyses. However, experimental validation will be required to confirm the biochemical activities and RNA-related functions of these proteins in *M. polymorpha*.

**Figure 10.**
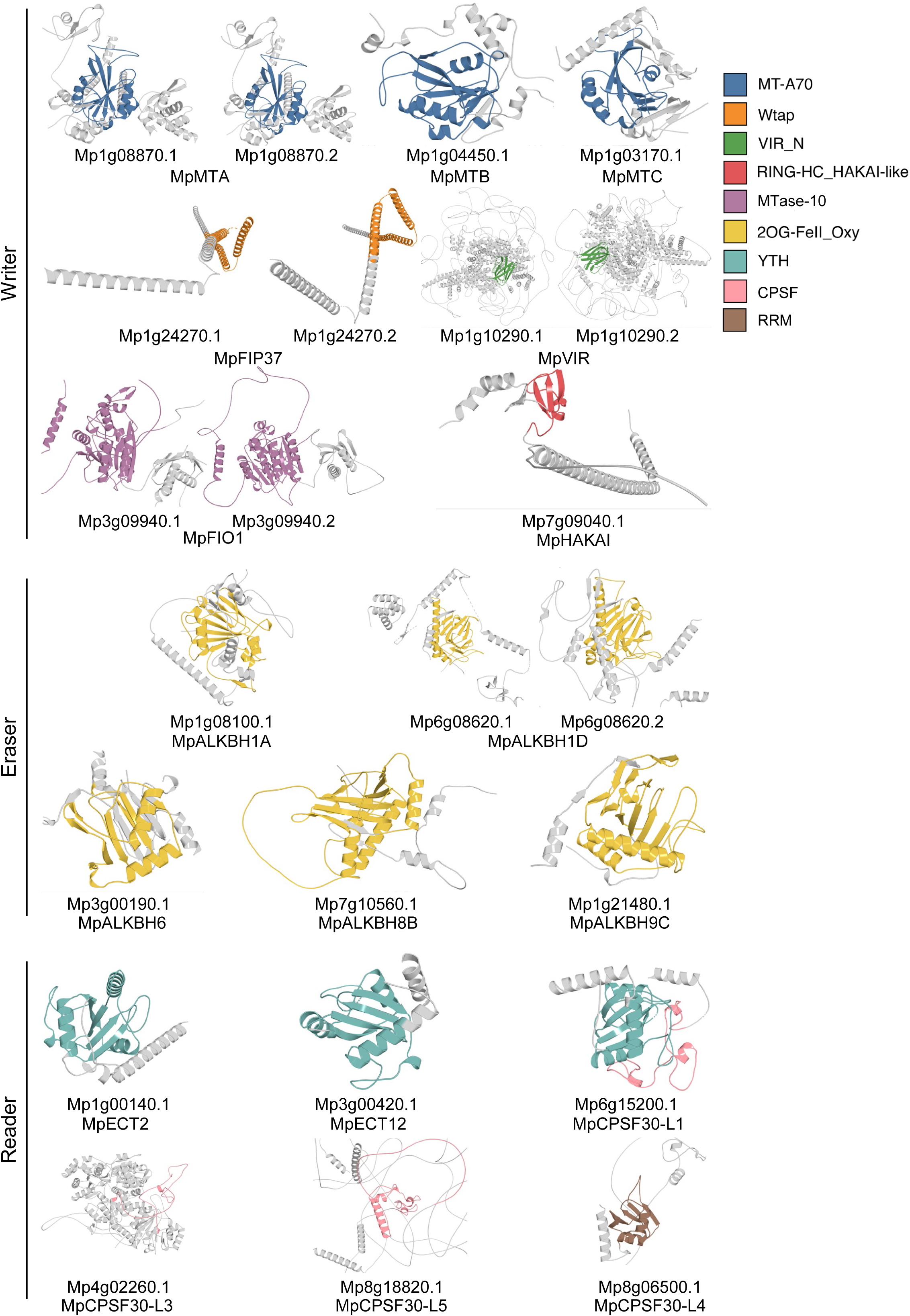
Predicted three-dimensional structures of *M. polymorpha* candidate m^6^A-related protein isoforms. Structures are grouped as writers, erasers, and readers, with protein identifiers and family names beneath the models. Models were obtained from the AlphaFold Protein Structure Database except for the two MpVIR isoforms, which were modeled using ColabFold. Colored ribbon segments indicate annotated domains: MT-A70 (blue), Wtap (orange), VIR_N (green), RING-HC_HAKAI-like (red), MTase-10 (mauve), ALKBH (yellow), YTH (teal), CPSF (pink), and RRM (brown). Displayed regions outside annotated domains are grey; low-confidence regions excluded for visualization are not shown.

### A predicted protein-association network links *M. polymorpha* m^6^A-related proteins to RNA-processing factors

Finally, we used STRING to examine predicted functional associations among the identified writer, eraser, and reader proteins and between these proteins and additional proteins (Figure 11). The network contained one main connected component and three isolated protein nodes labelled MpALKBH1D, MpCPSF30-L3, and MpCPSF30-L5. Most candidate m^6^A-related proteins were incorporated into the main component, although their connectivity and associated partners differed among the three functional groups.

**Figure 11.**
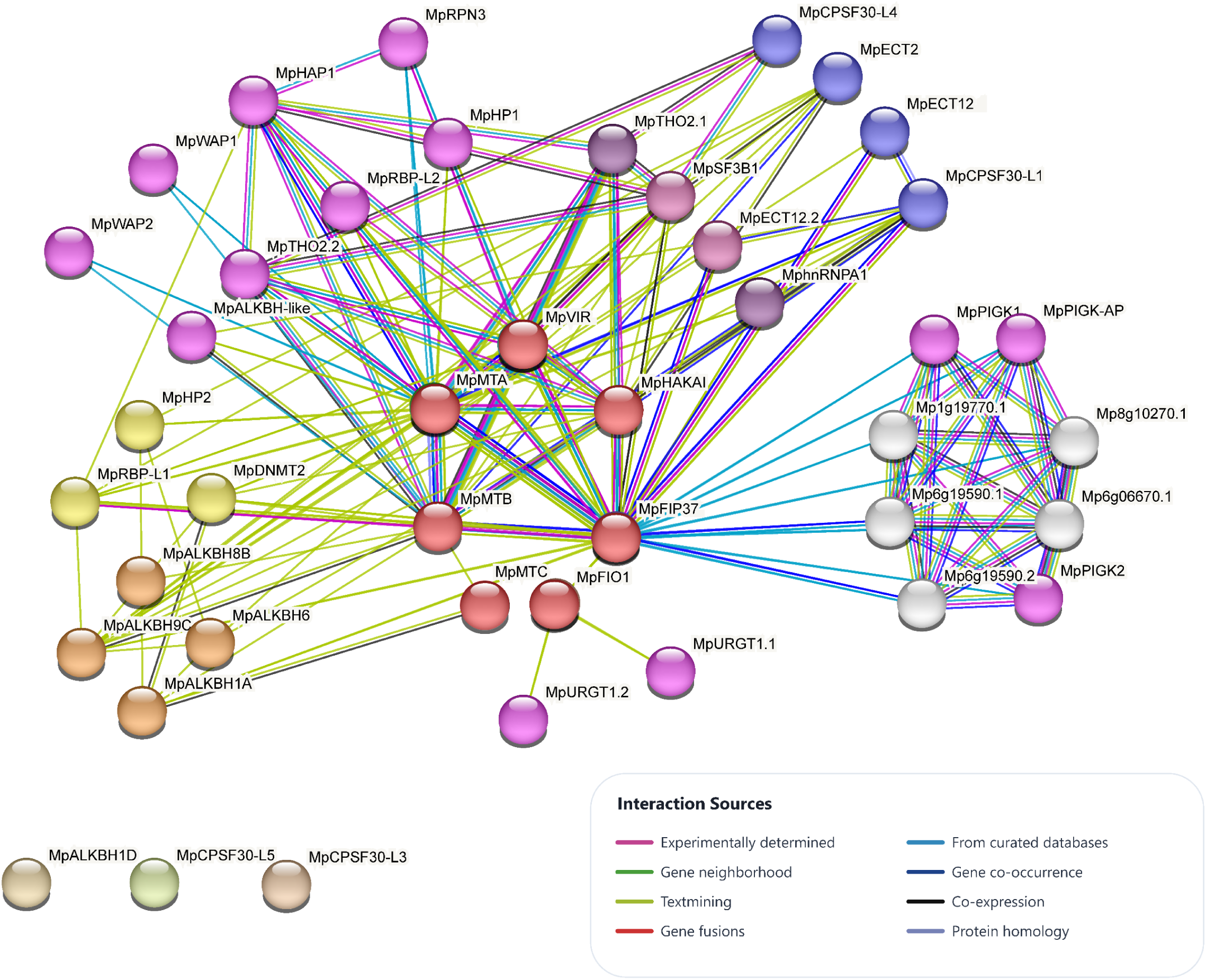
STRING-derived protein-protein interaction (PPI) network of *M. polymorpha* candidate m^6^A-related proteins and predicted functional associates. Spheres denote labeled proteins. The 18 candidate query proteins were represented by one protein per locus; the .1 isoform was used for loci with multiple annotated isoforms. Within the main connected component, candidate writer nodes are salmon, eraser nodes orange, and reader nodes blue. The figure displays three unconnected candidate nodes - MpALKBH1D, MpCPSF30-L3, and MpCPSF30-L5 - below the connected component. Edge colors indicate STRING evidence channels: experimentally determined (magenta), gene neighborhood (green), text mining (olive), gene fusion (red), curated databases (blue), gene co-occurrence (dark teal), co-expression (black), and protein homology (lavender). The minimum required STRING association score was 0.4, with up to 20 first-shell and five second-shell additional interactors. Edges represent predicted functional associations.

The writer-associated portion of the main component included MpMTA, MpMTB, MpMTC, MpFIP37, MpVIR, MpFIO1, and MpHAKAI. Within this component, MpMTA, MpMTB, MpFIP37, MpVIR, and MpHAKAI showed predicted associations with other writer candidates and additional proteins. Several associated proteins were related to RNA binding or RNA processing, including MpRBP-L1, MpRBP-L2, MpTHO2.1, MpTHO2.2, MpSF3B1, and MphnRNPA1. Studies in other species have implicated THO/TREX components in RNA processing and export and established roles for SF3B1 and hnRNP A1 in pre-mRNA splicing (Butt et al., 2021; Francisco-Mangilet et al., 2015; Tavanez et al., 2012). Their occurrence within the writer-associated region therefore suggests potential functional connections between the candidate m^6^A machinery and broader RNA-processing pathways. MpFIP37 was also connected to a peripheral subnetwork containing MpPIGK1, MpPIGK2, MpPIGK-AP, and several proteins labeled with locus identifiers. Reader candidates, including MpECT2, MpECT12, MpCPSF30-L1, and MpCPSF30-L4, were also included in the main connected component. Their predicted associations with writer candidates and proteins annotated as RNA-processing factors suggest possible links between the reader candidates and RNA-processing pathways. However, MpCPSF30-L3 and MpCPSF30-L5 appeared as isolated nodes under the network settings used in this analysis.

Among the eraser candidates, MpALKBH1A, MpALKBH6, MpALKBH8B, and MpALKBH9C were incorporated into the main component, whereas the MpALKBH1D node was isolated. The connected ALKBH candidates showed predicted associations with one another and with additional proteins annotated as RNA-related factors, suggesting possible links to RNA metabolism. The absence of predicted edges for MpALKBH1D, MpCPSF30-L3, and MpCPSF30-L5 under the selected network parameters does not demonstrate the absence of biological interactions, because such interactions may not yet be represented in the databases or may occur only under specific developmental or environmental conditions.

The network was inferred using STRING’s integrated functional-association evidence and does not establish direct physical interactions in *M. polymorpha* (Szklarczyk et al., 2023). Collectively, the predicted network links subsets of the *M. polymorpha* candidate m^6^A-related proteins to additional proteins annotated as RNA-related factors. These predictions identify candidate associations for subsequent biochemical and genetic testing.

## Discussion

Our analysis identifies 18 candidate m^6^A-related loci in *Marchantia polymorpha* and places them in a comparative framework spanning 119 plant genomes. The candidates encompass seven writer-related proteins, five ALKBH-family proteins, and six YTH- or CPSF30-related proteins, represented by 23 selected transcript models. Their phylogenetic placement and domain composition support the proposed family assignments, while the developmental and stress-expression profiles indicate that the corresponding genes are regulated differently rather than expressed as a single unit (Figures 2, 3, and 6-9). These data provide a framework for studying RNA methylation in *M. polymorpha*.

The detection of MpMTA, MpMTB, MpFIP37, MpVIR, and MpHAKAI is consistent with the conservation of the core mRNA writer machinery described in *Arabidopsis thaliana* (Figures 2 and 3) (Ruzicka et al., 2017; Zhong et al., 2008). This finding extends earlier reports that recognized several writer homologs in *M. polymorpha*, while the present study analyzes them alongside eraser and reader-related candidates (Pietrykowska et al., 2022). MpMTC and MpFIO1 warrant separate consideration. Although MpMTC contains an MT-A70-related domain, its participation in the canonical writer complex cannot be assumed from that feature. *A. thaliana* FIO1 modifies U6 snRNA and affects splice-site selection, so the annotation of MpFIO1 as writer-related does not establish that it acts on the same RNAs as MpMTA (Parker et al., 2022). Across the sampled species, relatively limited copy-number variation among several writer families contrasts with broader variation in ALKBH- and reader-related groups (Figure 2). Family counts reported for flowering plants and gymnosperms show a similar overall diversity, although direct numerical comparisons depend on genome assembly, isoform filtering, and family definitions (Hasan et al., 2024; Yao et al., 2024; Zhang et al., 2024).

The eraser and reader-related candidates illustrate why domain-based annotation must be interpreted at the family level. All five MpALKBH proteins contain an ALKBH-related domain, but the present data cannot show that each removes m^6^A from mRNA (Duan et al., 2017; Leihne et al., 2011; Martinez-Perez et al., 2017). Likewise, the YTH domains of MpECT2 and MpECT12 provide a rationale for testing m^6^A recognition, whereas the CPSF30-like proteins are not all equivalent to the experimentally characterized *A. thaliana* CPSF30-L reader (Arribas-Hernandez et al., 2018; Song et al., 2021). In particular, MpCPSF30-L3, MpCPSF30-L4, and MpCPSF30-L5 lack a detectable YTH domain in the displayed annotation (Figure 3B). They should therefore remain reader-related candidates until m^6^A binding is demonstrated. The predicted structures provide structural context for the sequence-based domain annotations but cannot resolve substrate specificity, catalytic activity, or RNA-binding capacity (Figure 10). Previous heterologous complementation by a *M. polymorpha* YTHDF-like protein supports conservation of some reader-associated molecular function; it does not establish the endogenous functions of the candidates analyzed here (Flores-Tellez et al., 2023).

Chromosomal comparisons add a separate layer of evolutionary evidence. Eight of the 18 *M. polymorpha* candidate loci had detectable collinear relationships with *Marchantia quadrata*, including members of each of the three functional groups (Figure 4). The other ten loci lacked a link under the selected criteria, which cannot be equated with the absence of a homolog in *M. quadrata*. The relationships indicate retention of local genomic context for a subset of candidates without establishing one-to-one orthology or the direction of any duplication or loss. In this respect, the collinearity analysis complements sequence phylogenies but does not replace locus-specific evolutionary analysis.

The putative promoter analysis suggests possible routes by which the candidate genes could respond to developmental and environmental signals. We detected 636 annotated motif occurrences across 23 upstream transcript entries representing 18 loci (Figure 5 and Table 2). Hormone-responsive motifs were the most abundant broad category, whereas the general light-responsive motif was the most frequent individual class. These counts describe predicted sites rather than 636 distinct regulatory events; transcript entries from the same locus are also not independent genes. Because the analyzed regions were defined relative to annotated transcript 5′ ends rather than experimentally mapped transcription start sites, motif identities and positions require experimental confirmation. Functional labels derived from seed-plant studies, such as endosperm- or seed-associated elements, should not be transferred literally to the liverwort life cycle.

The expression analyses indicate potential developmental contexts in which these genes could be investigated. Public expression data varied across the sporeling time course, vegetative tissues, reproductive structures, and sporophyte samples (Figure 6). In the independent qRT-PCR experiment, the thallus showed higher transcript abundance than gemmae and gemma cups for many candidates, including MpMTA, MpMTC, MpFIP37, MpVIR, MpFIO1, all readers, and all CPSF30-like genes (Figure 7). Not all genes followed the same pattern: MpMTB transcript abundance in gemma cups was comparable to that in the thallus or gemmae cup, and MpECT12 abundance in gemma cups exceeded that in gemmae while remaining below that in the thallus. These differences are consistent with regulated deployment of RNA-related factors across vegetative and asexual tissues, but transcript abundance alone cannot establish a developmental function or a change in m^6^A deposition. The broader database profiles and the qRT-PCR experiments sample different developmental materials and should be interpreted as complementary views rather than paired measurements of the same specimens.

Abiotic stress also elicited gene-specific responses. Reanalysis of the public RNA-seq data detected changes under single and combined stresses, with some combinations differing from the corresponding individual treatments (Figure 8). In our qRT-PCR experiments, darkness was associated with increased transcript abundance for several candidates, including MpMTA, MpMTC, MpVIR, MpFIO1, MpALKBH1A, MpECT2, MpCPSF30-L1, and MpCPSF30-L3; cold, heat, osmotic treatment, and salinity produced different responses across the gene set (Figure 9). This diversity resembles the context-dependent expression reported in other plant gene-family surveys, but the RNA-seq and qRT-PCR datasets differ in treatment design and biological material (Hasan et al., 2024; Shan et al., 2025; Yao et al., 2024). More importantly, altered expression of a candidate regulator is not a direct measure of cellular m^6^A abundance, target-RNA modification, or stress tolerance. Time-resolved methylation and functional assays will be needed to determine whether any of these transcriptional responses changes RNA fate during stress.

The predicted association network provides possible connections to broader RNA metabolism. Several writer-related candidates appear within the main component together with proteins annotated for RNA binding, processing, or export, whereas MpALKBH1D and selected CPSF30-like candidates lacked edges under the selected network settings (Figure 11). STRING evidence supports predicted functional associations, which cannot be interpreted as proof of direct physical interactions (Szklarczyk et al., 2023). Predicted cellular localizations also warrant direct testing: for example, the nuclear localization predicted for MpECT2 should be examined in light of the cytoplasmic localization reported for *A. thaliana* ECT2, ECT3, and ECT4 (Arribas-Hernandez et al., 2018). Localization imaging, biochemical interaction tests, and RNA-binding assays would distinguish these possibilities.

The combined results prioritize experimentally tractable questions. Loss-of-function and complementation analyses could test whether specific candidate genes affect gemma cup development or stress responses; m^6^A mapping and measurements on defined RNA classes could then establish which effects depend on methylation. Direct assays of MpMTA- and MpFIO1-related activities, ALKBH-dependent demethylation, YTH-dependent RNA binding, and CPSF30-associated 3′-end processing would clarify whether the functions described in flowering plants are retained or altered in *M. polymorpha* (Arribas-Hernandez et al., 2018; Duan et al., 2017; Parker et al., 2022; Ruzicka et al., 2017; Song et al., 2021). Such experiments are necessary to move from a conserved and transcriptionally responsive candidate repertoire to an experimentally supported model of liverwort RNA methylation.

## Conclusion

This study identifies 18 candidate m^6^A-related genes in *Marchantia polymorpha* - seven writer-related, five ALKBH-related, and six reader or reader-related loci - and places them within a survey of 119 plant genomes. Conserved domains and phylogenetic relationships support their family assignments, while collinearity with *Marchantia quadrata* identifies eight candidates with retained genomic context. Promoter analysis identifies putative regulatory elements, while expression analyses reveal variation across tissues, developmental stages, and abiotic stress conditions. Predicted protein structures and associations provide additional context for investigating the functions of the identified proteins. Together, these findings provide a comparative framework for testing how RNA methylation is established, interpreted, and regulated in liverworts. Direct measurement of RNA modifications, biochemical activity, and genetic phenotypes remains essential to determine the functions of individual candidates.

---

**Supplementary Figure 1.**
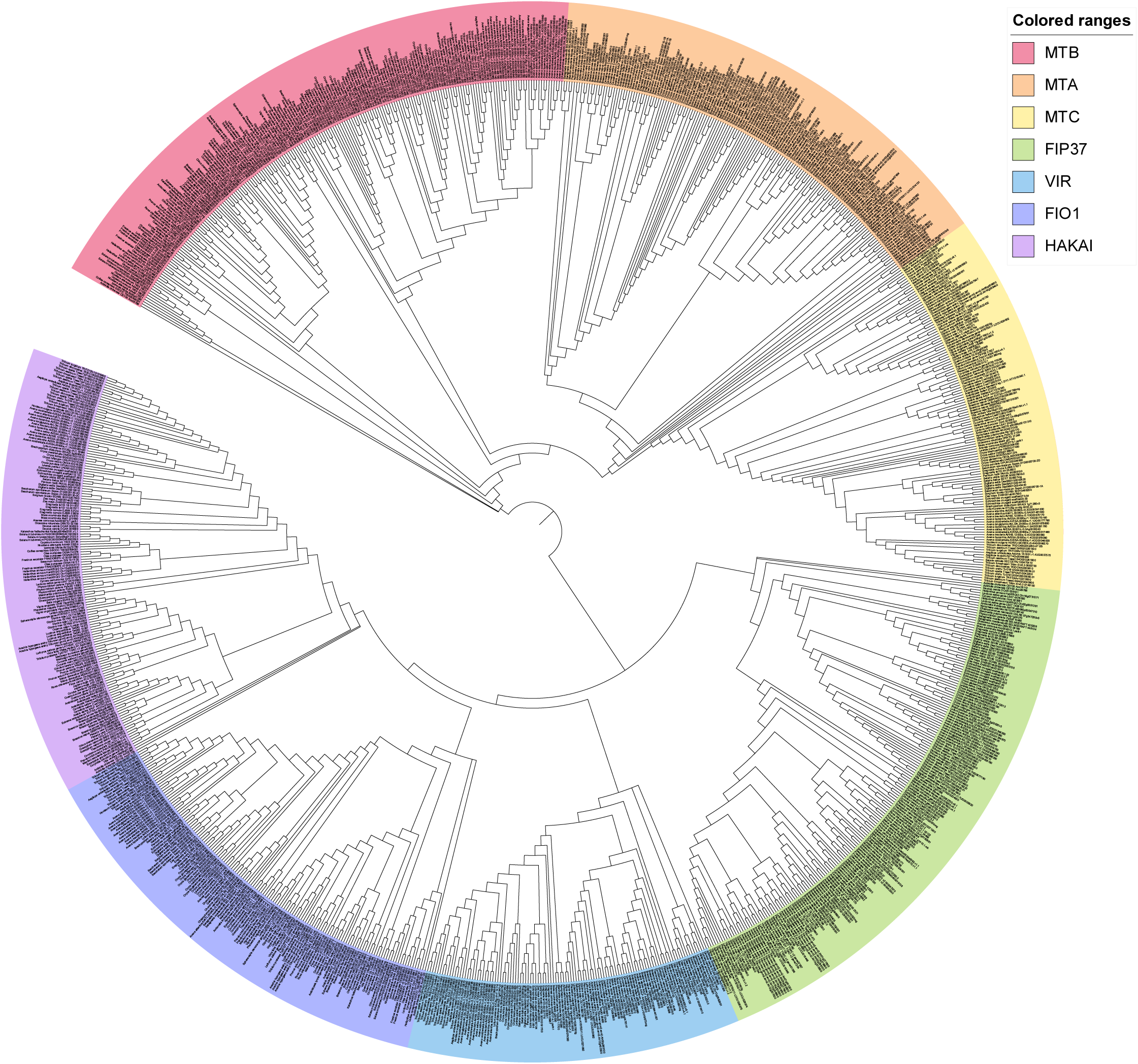
Composite display of maximum-likelihood phylogenies of candidate m^6^A writer proteins across 119 plant species. Colored outer sectors delimit MTB (pink), MTA (peach), MTC (pale yellow), FIP37 (pale green), VIR (light blue), FIO1 (periwinkle), and HAKAI (lavender) clades. Within each domain-defined writer group, branches represent IQ-TREE-inferred protein relationships. The arrangement of the writer groups was imposed using a synthetic backbone for visualization and does not represent an inferred relationship among the groups; bootstrap support therefore applies only within each domain-defined subtree. One representative isoform per locus was retained.

**Supplementary Figure 2.**
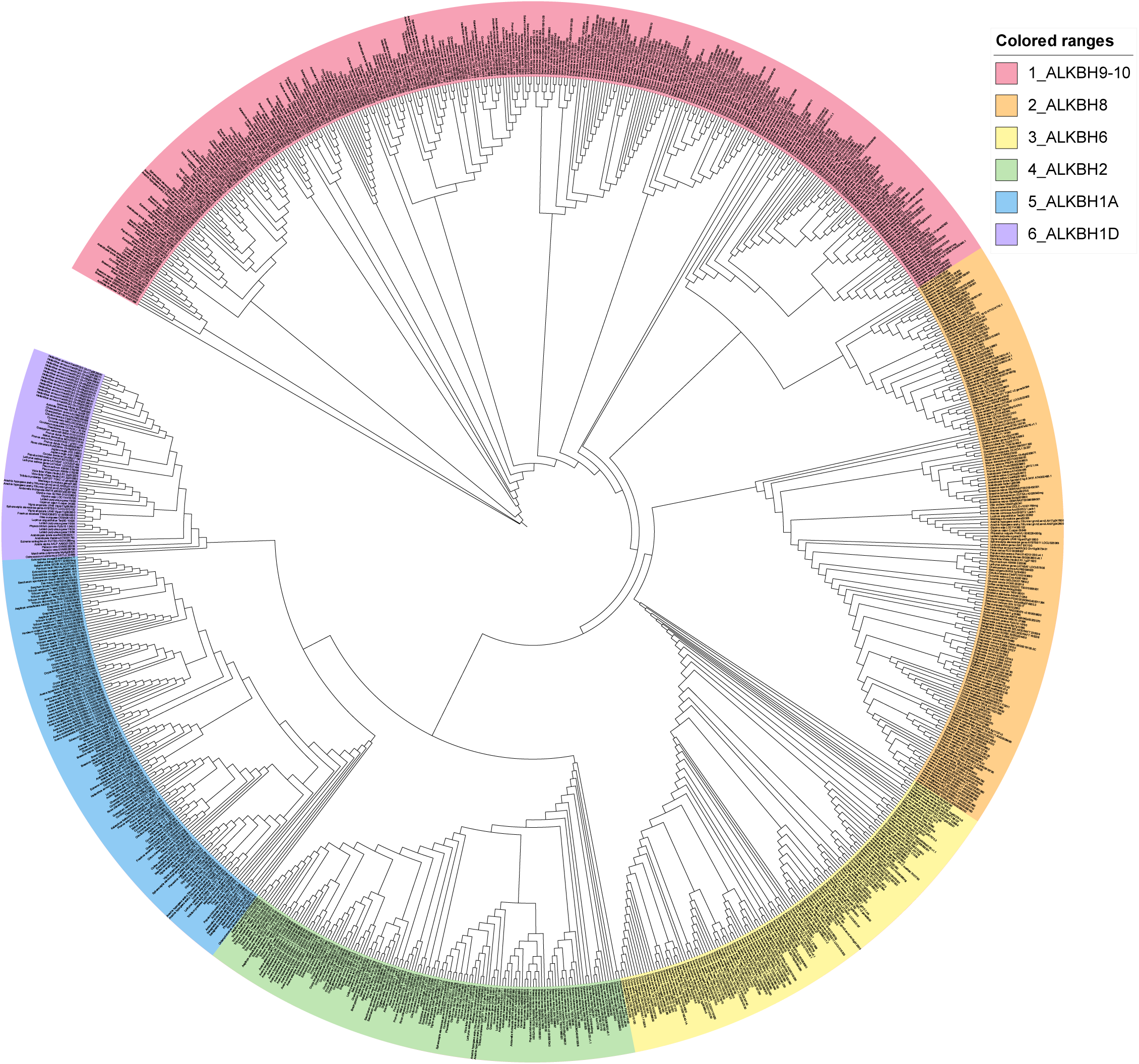
Maximum-likelihood phylogeny of candidate ALKBH proteins across 119 plant species. Colored outer sectors identify ALKBH9/10 (pink), ALKBH8 (peach), ALKBH6 (pale yellow), ALKBH2 (pale green), ALKBH1A (light blue), and ALKBH1D (lavender) groups. Numeric prefixes in the key identify groups, rather than gene copy numbers. One representative isoform per locus was retained.

**Supplementary Figure 3.**
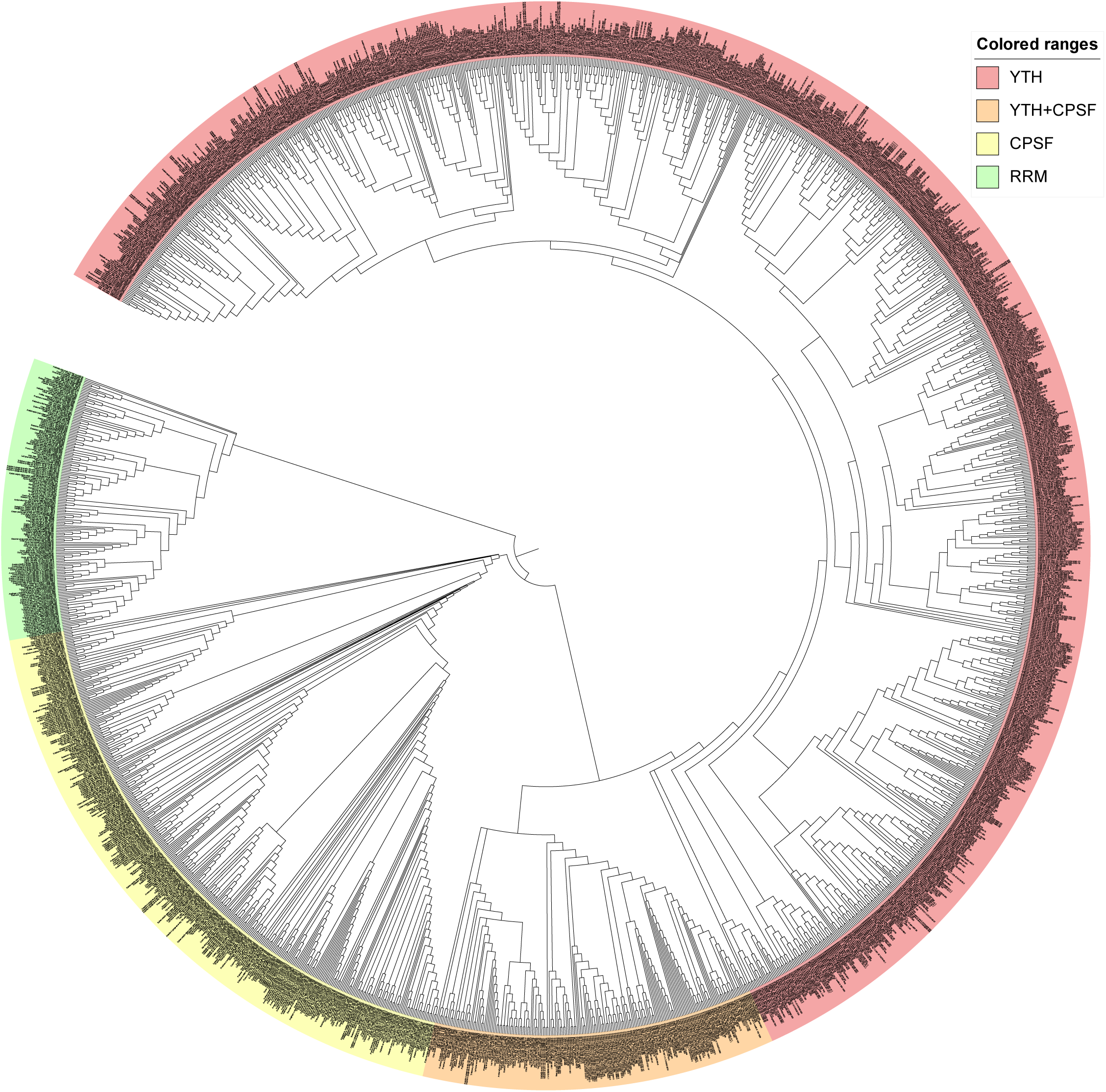
Composite display of maximum-likelihood phylogenies of candidate m^6^A reader-related proteins across 119 plant species. Colored outer sectors distinguish YTH (pink), combined YTH+CPSF (peach), CPSF (pale yellow), and RRM (pale green) protein/domain categories. Within each domain-defined reader group, branches represent IQ-TREE-inferred protein relationships. The arrangement of the reader groups was imposed using a synthetic backbone for visualization and does not represent an inferred relationship among the groups; bootstrap support therefore applies only within each domain-defined subtree. One representative isoform per locus was retained.

**Supplementary Figure 4.**
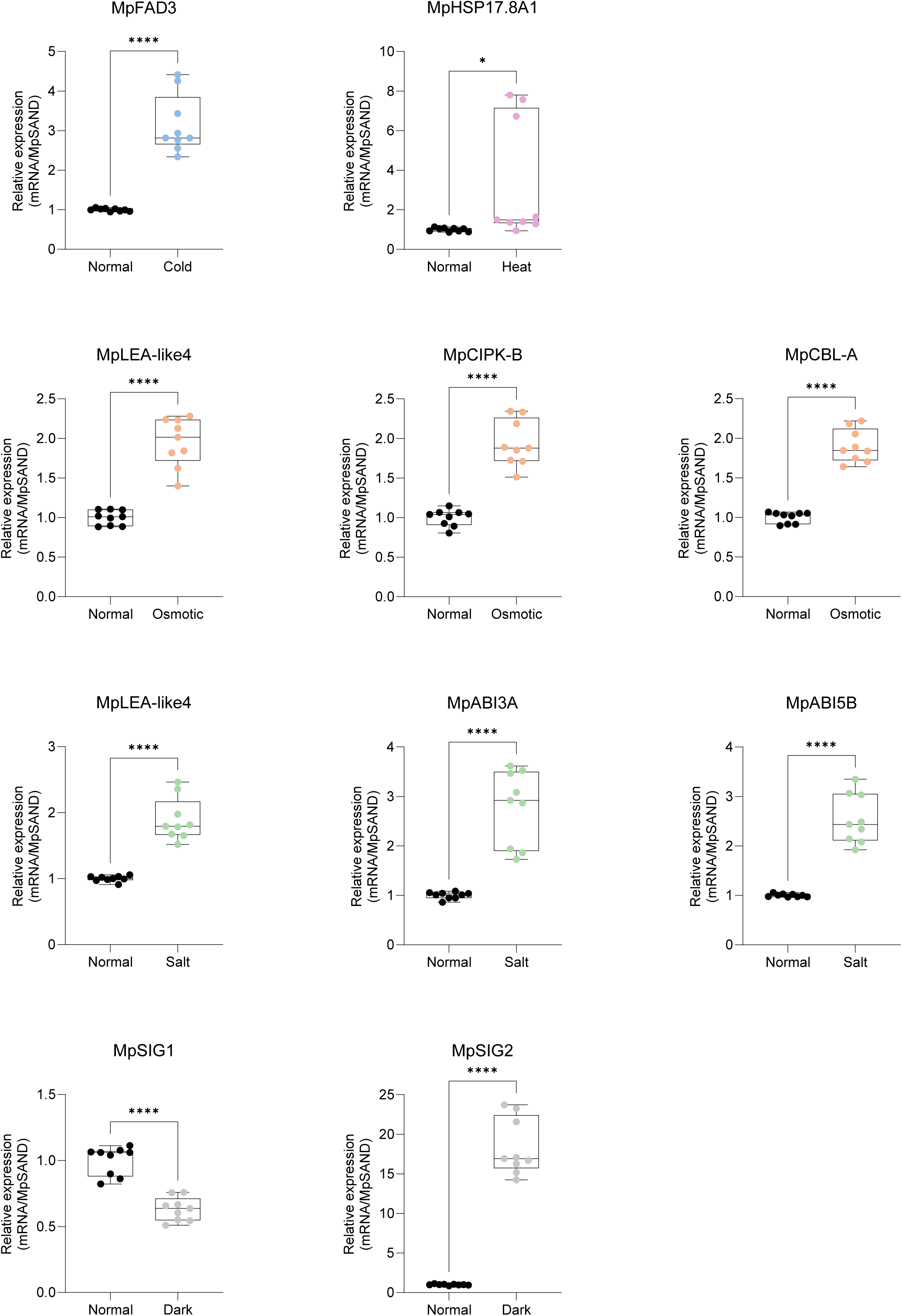
qRT-PCR expression of abiotic-stress marker genes under the treatments used in this study. *MpFAD3* is shown under cold; *MpHSP17.8A1* under heat; *MpLEA-like4*, *MpCIPK-B*, and *MpCBL-A* under osmotic stress; *MpLEA-like4*, *MpABI3A*, and *MpABI5B* under salt; and *MpSIG1* and *MpSIG2* under darkness. Each treatment is compared with its Normal control. *MpSAND* was the expression reference. Control points are black; cold, heat, osmotic, salt, and dark-treatment points are blue, pink, orange, green, and grey, respectively. Boxes span the first to third quartiles, center lines indicate medians, whiskers indicate the plotted ranges, and points show individual measurements. Brackets indicate comparisons assessed by two-tailed Welch’s *t* test; * indicates *P* < 0.05 and **** indicates *P* < 0.0001.

## Notes

### Competing Interest Statement

The authors have declared no competing interest.

### Funding

- Suzhou Science and Technology Bureau: ZSL2024334
- Startup Fund, Duke Kunshan University: 00AKUG0122

